# The Genomic portrait of the Picene culture: new insights into the Italic Iron Age and the legacy of the Roman expansion in Central Italy

**DOI:** 10.1101/2024.03.18.585512

**Authors:** Francesco Ravasini, Helja Niinemäe, Anu Solnik, Luciana de Gennaro, Francesco Montinaro, Ruoyun Hui, Chiara Delpino, Stefano Finocchi, Pierluigi Giroldini, Oscar Mei, Elisabetta Cilli, Mogge Hajiesmaeil, Letizia Pistacchia, Flavia Risi, Chiara Giacometti, Christiana Lyn Scheib, Kristiina Tambets, Mait Metspalu, Fulvio Cruciani, Eugenia D’Atanasio, Beniamino Trombetta

## Abstract

**Background:** The Italic Iron Age was characterized by the presence of various ethnic groups partially examined from a genomic perspective. To explore the evolution of Iron Age Italic populations and the genetic impact of Romanization, we focused on the Picenes, one of the most fascinating pre-Roman civilizations, who flourished on the Middle Adriatic side of Central Italy between the 9^th^ and the 3^rd^ century BCE, until the Roman colonization.

**Results:** We analyzed more than 50 samples, spanning more than 1,000 years of history from the Iron Age to Late Antiquity. Despite cultural diversity, our analysis reveals no major differences between the Picenes and other coeval populations, suggesting a shared genetic history of the Central Italian Iron Age ethnic groups. Nevertheless, a slight genetic differentiation between populations along the Adriatic and Tyrrhenian coasts can be observed, possibly due to genetic contacts between populations residing on the Italian and Balkan shores of the Adriatic Sea. Additionally, we found several individuals with ancestries deviating from their general population. Lastly, In the Late Antiquity period, the genetic landscape of the Middle Adriatic region drastically changed, indicating a relevant influx from the Near East.

**Conclusions:** Our findings, consistently with archeological hypotheses, suggest genetic interactions across the Adriatic Sea during the Bronze/Iron Age and a high level of individual mobility typical of cosmopolitan societies. Finally, we highlighted the role of the Roman Empire in shaping genetic and phenotypic changes that greatly impacted the Italian peninsula.

## Background

Before the unification under Roman rule, the inhabitants of the Italian peninsula consisted of various regional groups characterized by specific geographical distribution, cultural identities, and languages [1–3]. The Italic IA (about 10^th^- 3^rd^ centuries BCE) was a transformative era characterized by the use of new metals in artifact production, novel agricultural and breeding practices, linguistic and cultural development, migrations and conflicts. During the Late Bronze Age, distinct Italic ethnicities emerged, giving rise to new communities with well-defined cultural and linguistic identities that consolidated during the IA, such as the Picenes, Etruscans, Latins, and others [1,4–6]. These cultural entities were connected to each other and engaged in close commercial and cultural contacts [1]. Moreover, the presence of non-autochthonous goods in Italic Iron Age archaeological sites suggests that these populations were part of an extensive commercial network, connecting the Italian Peninsula with the whole Mediterranean basin and Europe [7].

Despite extensive archaeological and historical studies, the genetic origins and possible admixture events among these populations are still elusive. The population dynamics that have contributed to shape the ancient and modern Italian gene pool remain largely unknown, and only a limited number of studies have investigated the genomic variability of Italic IA ethnicities [8–10]. In this context, one of the understudied areas is the Adriatic side of central Italy, nowadays the Marche region. In this area, between the 9^th^ and the 3^rd^ century BCE one of the most flourishing pre-Roman civilizations was established: the Picenes [1,5,11–13].

This term refers more to an ethnicity rather than a population. Indeed, the Picenes were divided into many local groups not necessarily ancestrally related, but sharing a common cultural substratum [5]. According to a mythical tale reported by Pliny the Elder (1^st^ century BCE), the Picenes are connected with the Sabines, from whom they would have separated in search of a new homeland. This migration occurred in the ritual form of a so-called sacred spring (*ver sacrum*) following a totemic animal, the woodpecker, in Latin *picus*. Thus, the migrants assumed the ethnonym of *Picenes* (or *Picentes*), i.e., "people of the woodpecker” [5]. Regardless of their traditional ethnogenesis, it has been hypothesized that the Picenes derive from Late Bronze Age cultures of the Adriatic coast with connections to other coeval ethnicities such as the trans-Adriatic ones [5].

Our current knowledge about this civilization is mainly based on archaeological findings belonging to grave goods found in necropolises. The funerary settings show a society organized into coherent (but differentiated) political groups rooted in local traditions but, at the same time, open towards foreign cultural inputs which were often converted into an original artistic conception [5,11,12]. In this context, it has been suggested that Picenes had frequent contacts with other cultures, for example with Northern Europe and Eastern Mediterranean populations, as traced by many artifacts found in the burials [5,13]. The relevance of these cultural contacts in shaping the gene pool of the area is still debated, especially since later events have probably blurred these genetic traces.

In the early years of the 3^rd^ century BCE, the Picene culture started to fade due to the Roman expansion [5]. The Romans have deeply impacted the history of Italy by contributing to different socio-cultural aspects and, possibly, to demography and genetics [8,14]. The rise of a multicultural Roman Empire changed the genetic landscape of the city of Rome, introducing a strong genetic component from the Near East that also lasted throughout the Late Antiquity [8]. Although this shift in the gene pool has also been observed outside the city of Rome post [9,15], it is still unclear how pervasive it was in the entire peninsula.

To shed light on all these aspects, we conducted an archaeogenetic analysis of the Italian Middle Adriatic populations covering a period of more than 1,000 years by performing shotgun sequencing of 102 ancient individuals. In particular, we analyzed: two different Picene necropolises (Novilara and Sirolo-Numana) both dated to the early stages of this civilization (8^th^-5^th^ centuries BCE); an Etruscan necropolis (Monteriggioni/Colle di Val d’Elsa, 8^th^-6^th^ centuries BCE) to compare the gene pool of the two side of central Italy during the IA; and finally, a Late Antiquity funerary site (Pesaro) dated to the 6^th^-7^th^ centuries CE and located 7 km away from Novilara, to study the genetic changes in the Picene territory after the Romanization (Figure 1, Additional file 1). With our study, we described the gene pool of the Picenes shedding light on their genetic origin, deeply connected to the other coeval populations. Nevertheless, we highlighted relevant differences between the IA populations settled in the Adriatic and the Tyrrhenian coasts of Italy. Finally, we assessed the genetic impact of the historical events related to Romanization in the former Picene area.

**Figure 1.**
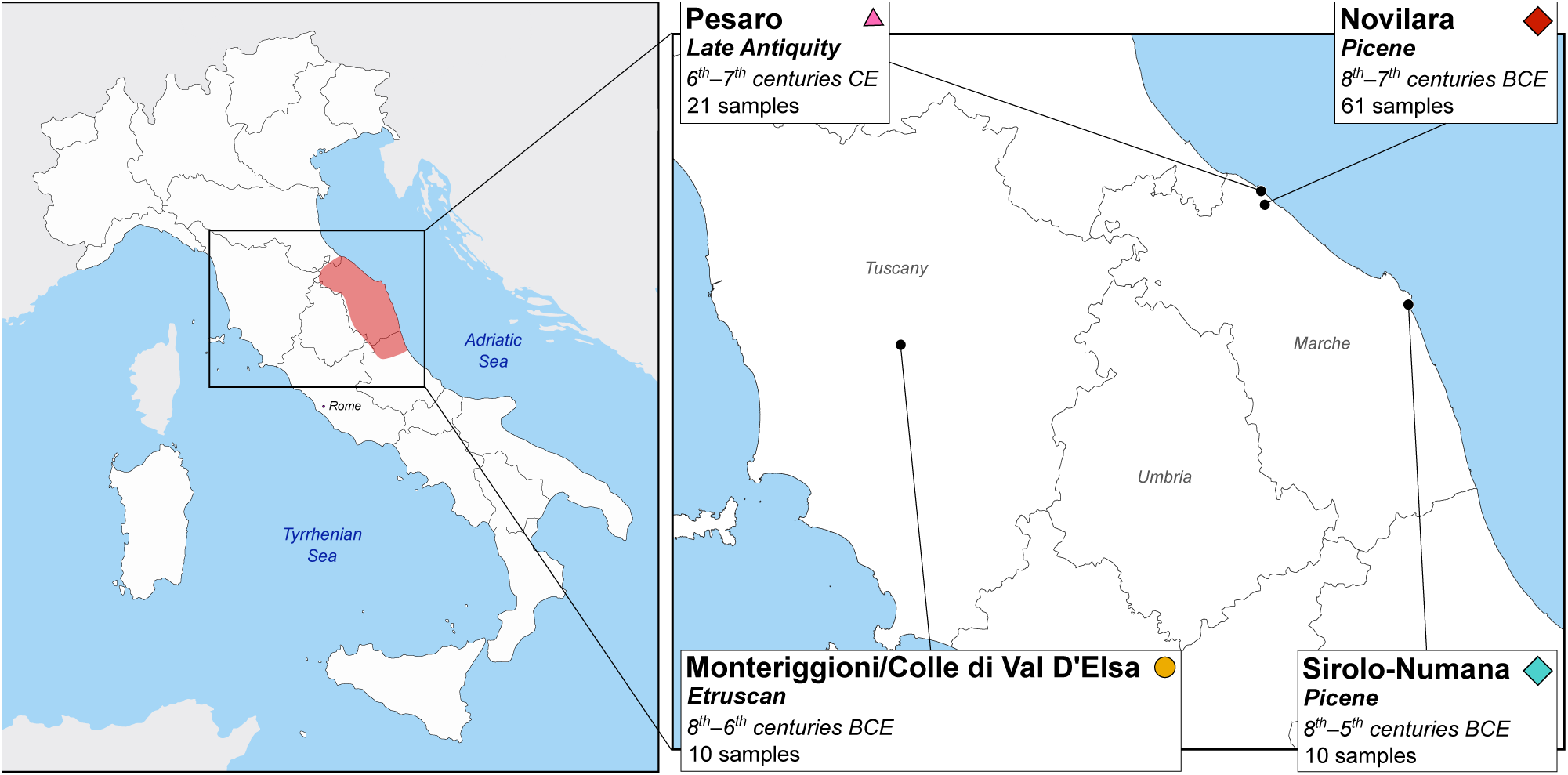
Location of the sites analyzed in this study. On the left, map of Italy with the Picene area highlighted in red. On the right, magnification of Central Italy showing the location, the period, and the number of samples for each necropolis analyzed in this study. Symbols associated with necropolises are the same as in PCA (Figure 2A).

## Results

We collected and extracted DNA from tooth and petrous bone samples of 71 Picene individuals (61 from Novilara necropolis and 10 from Sirolo-Numana necropolis), 10 Etruscans from Monteriggioni/Colle di Val d’Elsa necropolis and 21 Late Antiquity individuals from Pesaro (hereafter called Pesaro Late Antiquity), resulting in a total of 102 samples (Fig 1; Additional file 2: Table S1). For each sample, we built libraries and performed shotgun whole genome sequencing (see Methods section for details). We restricted all subsequent genomic analysis to individuals that passed authentication and quality control, for a total of 55 samples, with a nuclear genome coverage spanning from 0.04x to 0.33x (mean 0.15x) (Additional file 2: Table S1). To identify genetically identical samples, we performed kinship analysis (Additional file 2: Table S2 and S3) finding two Etruscan samples from Monteriggioni/Colle di Val D’Elsa (EV7A and EV19) possibly representing twins or the same individual. Therefore, we removed the one with the lowest coverage (EV19) for a final sample set of 54 individuals (Table 1).

**Table 1.** Samples analyzed, included in genetic population analysis and genomic imputation for each site. *Samples used for population genome-wide analysis. **Samples used for genomic imputation and haplotype-based analysis.

| Archaeological site | Group | Nr | Coverage $\geq 0.04X^*$ | Coverage $\geq 0.1X^{**}$ |
| --- | --- | --- | --- | --- |
| Novilara | Iron Age/Picene | 61 | 40 | 28 |
| Sirolo-Numana | Iron Age/Picene | 10 | 3 | 2 |
| Monteriggioni/Colle di Val d'Elsa | Iron Age/Etruscan | 10 | 4 | 1 |
| Pesaro | Late Antiquity | 21 | 7 | 4 |
|  | **Total** | **102** | **54** | **35** |

In order to contextualize our ancient Italian sequences in the genetic variability of ancient and modern Eurasia and North Africa, we combined the genomic information of the newly genotyped samples with relevant individuals from the Allen Ancient DNA Resource (AADR, v. 52.2 [16]) (Additional file 2: Table S4 and S5).

### The Picene and the genomic landscape of the Italic Iron Age

We placed the genetic variability of the Picenes in the context of a European and Mediterranean framework through Principal Component Analysis (PCA), including a total of 1464 ancient and modern individuals (Fig. 2A; Additional file 3: Fig. S1). The Picene individuals constitute a wide cluster between the genetic variability of modern Italian, Balkan and Northern European populations. Remarkably, no major differentiation between individuals from the two Picene necropolises (Novilara and Sirolo-Numana) can be observed, pointing to the genetic similarity among the two Picene sites, notwithstanding differences in their material cultures [5,13]. In agreement with previous findings about other Italic Iron Age cultural groups [10], the Picenes show a slight deviation from the genetic distribution of modern Central Italians, being shifted towards Northern Italians and, more in general, Central Europeans (Additional file 3: Fig. S2). Unsupervised Admixture analysis [17] performed with 1708 modern and ancient Eurasian and North African individuals (Fig. 2B; Additional file 3: Fig. S3) supports the general genetic homogeneity observed among the two different Picene sites. The principal genetic ancestry for the two Picene groups is composed of Anatolian Neolithic and Eastern Hunter Gatherer (EHG)/Yamnaya (also referred to as Steppe BA) components (about 90%), with only minor proportions of Serbia Iron Gates Mesolithic and Caucasus Hunter Gatherer (CHG), similarly to other Italic IA groups (Fig. 2B; Additional file 3: Fig. S3).

**Figure 2.**
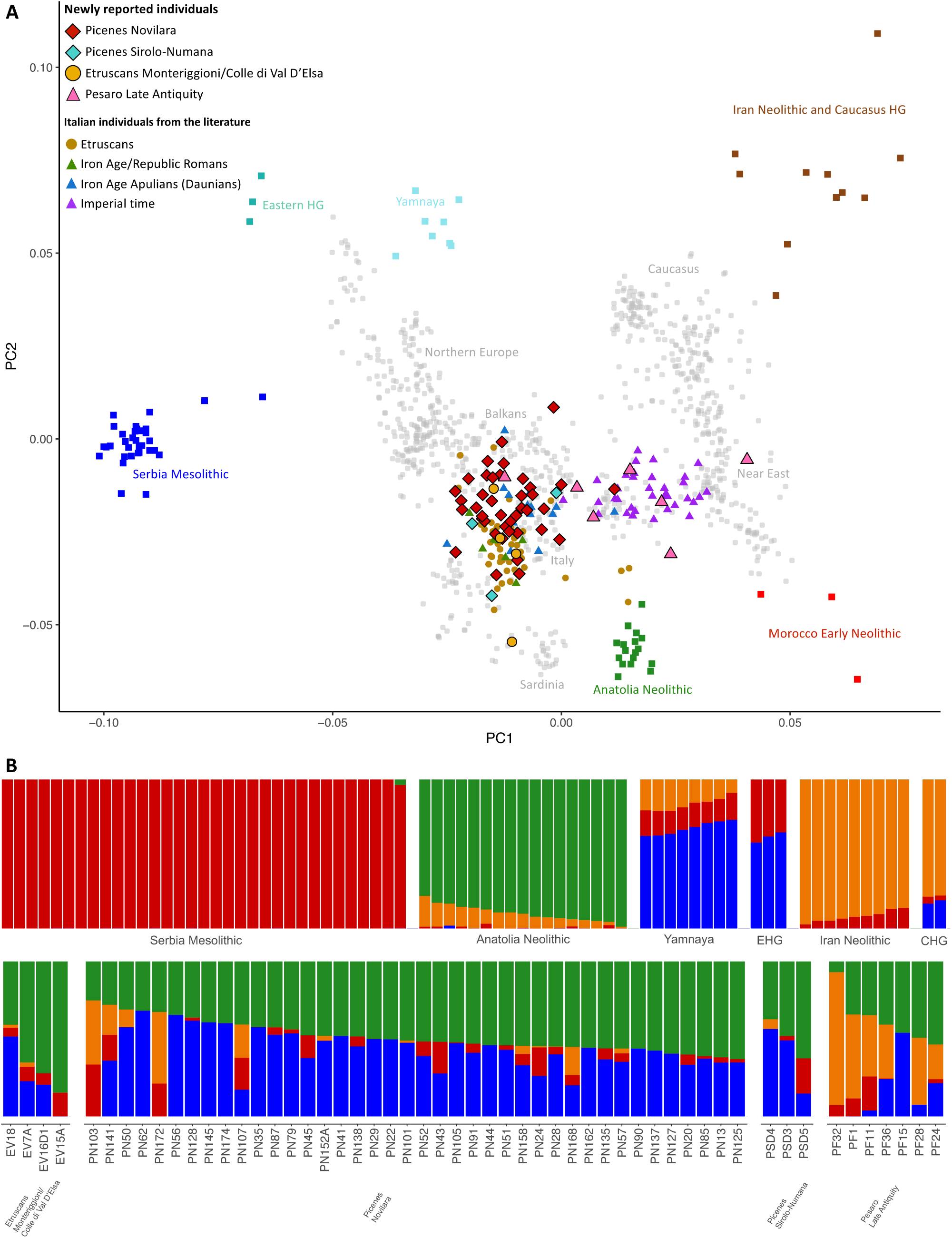
Population structure of the Italic IA and LA period. A) PCA with the newly reported samples and a relevant subset of modern and ancient individuals from the literature (see also Additional file 3: Fig. S1). Modern samples are pictured in gray. B) Unsupervised Admixture analysis at k=4. Above, a subset of samples representing the four ancestral components that contributed to the European gene pool since the Bronze Age: Serbia Iron Gates Mesolithic (red), Anatolian Neolithic (green), Caucasus Hunter-Gatherers (CHG)/Iran Neolithic (orange), and Eastern Hunter-Gatherers (EHG)/Yamnaya (blue); below, genetic make-up of the newly reported individuals.

To clarify the evolution of the gene pool of the Picenes and to get a general overview of the Italian IA population dynamics we also included in the downstream analyses newly generated data for 4 individuals from the Etruscan necropolis of Monteriggioni/Colle di Val d’Elsa (Table 1; Additional file 2: Table S1) together with Italian IA individuals from the literature [8–10]. The Monteriggioni/Colle di Val d’Elsa samples overlap the genetic variability of other Etruscan individuals [9] and mirror the ancestry patterns already identified in this ethnicity (Fig. 2; Additional file 3: Fig. S1 and S3).

Despite cultural differences, the PCA shows that IA populations exhibit relative genetic homogeneity, suggesting a shared genetic origin for these ethnicities in continuity with the former Italian Bronze Age (BA) cultures (Fig. 2A; Additional file 3: Fig. S1) [18]. Nevertheless, in the context of this genetic homogeneity, we can observe some differences, being the Picenes slightly shifted towards Balkan and Northern European modern populations. More in general, a significant differentiation is observed in the distribution along the PC2 of the Tyrrhenian people (Italy_IA_Romans and Etruscans, which are slightly closer to modern Sardinians) and the Adriatic group (Picenes and Italy_IA_Apulia, which tends to form a cluster shifted towards the ancient Yamnaya pastoralists) (Mann-Whitney U Test *p*-value < 0.0001; Additional file 3: Fig. S4). It may be possible that the Picenes (and more generally the IA Adriatic populations) have had slightly different evolutionary trajectories compared to the Tyrrhenian populations. This is confirmed by the D-statistics in the form D(X, Y; Italy_BA_EBA, YRI.SG) where X and Y are the Italic Iron Age groups and Italy_BA_EBA represents the previous Italian BA cluster. When comparing Italy_IA_Apulia and Picenes (the Adriatic groups), it is possible to observe a symmetrical relationship to the Italian Bronze Age (Fig. 3A; Additional file 2: Table S6). In the same way Etruscans and Italy_IA_Romans (the Tyrrhenian groups) are equally related to the previous Italian gene pool. On the other hand, the comparison of Tyrrhenian and Adriatic groups shows that the Bronze Age cluster tends to be more similar to the Tyrrhenian populations (Additional file 2: Table S6). Moreover, by performing D-statistics in the form D(Italy_IA_Adriatic, Italy_IA_Tyrrhenian; Italy_BA_EBA, YRI.SG), the Italian BA cluster results to be more related to the Tyrrhenian side (Fig. 3A; Additional file 2: Table S6). This result may suggest that the Adriatic and the Tyrrhenian sides of Italy were shaped by different demographic events after or during the Bronze Age.

**Figure 3.**
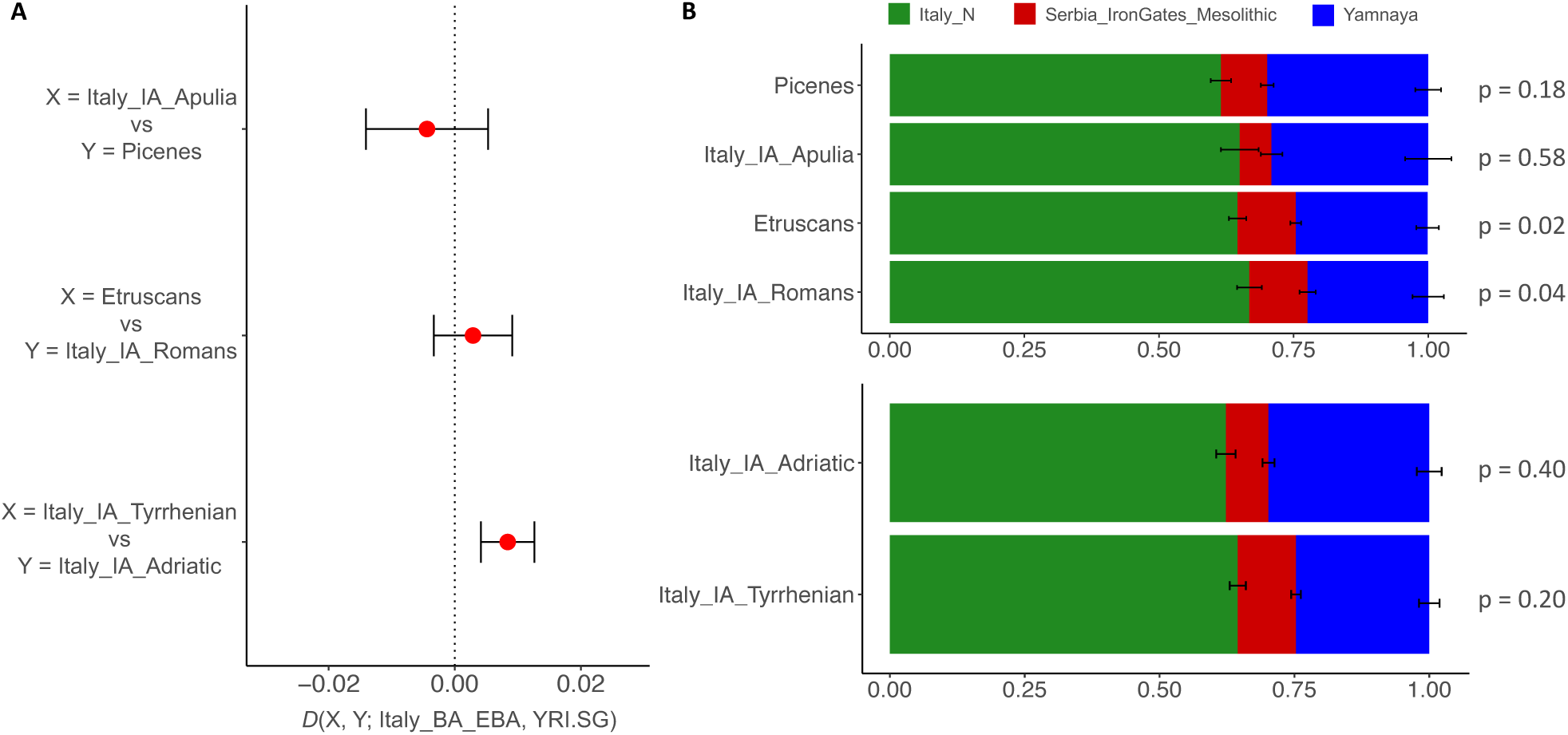
Genetic affinity and modeling of the Italic IA populations. A) D-statistics in the form D(X,Y; Italy_BA_EBA; YRI.SG), where X and Y are Italic IA groups. If D > 0, the Italian BA individuals (Italy_BA_EBA) are more closely related to population X, while if D < 0, Italy_BA_EBA is more closely related to population Y. In the first two tests, comparisons between Italic groups on the same side of the peninsula (Italy_IA_Apulia vs Picenes; Etruscans vs Italy_IA_Romans) are represented; the third test shows the comparison between all the Tyrrhenian vs all the Adriatic IA populations. B) qpAdm with 3 source populations for the Italic IA groups. Above, Italic IA ethnic groups analyzed so far. Below, Italic IA Adriatic and Tyrrhenian populations grouped together. For each model *p*-values are reported.

To explore which genetic influx may have primarily contributed to the evolution of the Adriatic gene pool we interpolated on a map the European ancestral components obtained with Admixture incorporating Eurasian individuals dating back to the 1^st^ Millennium BCE. The results reveal a nearly complementary distribution between the Anatolian Neolithic and EHG/Yamnaya components (Additional file 2: Table S7; Additional file 3: Fig. S5). Regions such as Etruria (i.e., the Etruscan region), Sardinia, and Southern Italy exhibit a higher prevalence of the Anatolian Neolithic component, while the Adriatic coast, particularly the Picene region, displays a more pronounced influence from the EHG/Yamnaya component (Additional file 3: Fig. S5).

To validate these findings, we analyzed the ancestry of these groups using a qpWave/qpAdm framework involving three source populations. When modeling these Iron Age (IA) groups with Serbia Iron Gates Mesolithic, Yamnaya, and Italy Neolithic (Italy_N) it emerges that the Adriatic people have a greater proportion of Yamnaya component compared to the Tyrrhenian ones (Fig 3B; Additional file 2: Table S8 and Table S9).

To look into the genetic relationships among the ancient Italian and European groups we performed a f4/NNLS analysis harnessing about 1.5 million of f4-statistics vectors as described in Saupe et al. [18]. We performed this analysis employing two different sets of ancestral components with the populations described in Lazaridis et al. [19] (Additional file 3: Fig. S6). The ancestral components that differ between the two sets are Yamnaya and EHG. We performed this analysis to potentially differentiate between these two genetic components, as they often appear similar when comparing Western European populations (as observed in Fig. 2B). Overall, this analysis recapitulates the emerging picture, highlighting the same ancestral components and with similar proportions for Bronze Age and Iron Age European populations, with the Adriatic populations showing a slightly greater Yamnaya component with respect to the Tyrrhenian groups. To have an overview of the phyletic relationships of the populations involved, we built an UPGMA tree based on the Euclidean distances between f4-statistics vectors. All the Italic IA ethnic groups are located in a single cluster suggesting a shared ancestral root (Additional file 3: Fig. S6). Notably, an Adriatic cluster emerges, encompassing Picenes and Italy_IA_Apulia, thereby distinguishing them from the Tyrrhenian populations. Remarkably, the sister clade to the IA of Italy is represented by trans-Adriatic cultures. This implies a potential influence from the Balkan peninsula on the evolution of the Italic gene pool, particularly impacting the Adriatic coasts of the peninsula.

The putative connection among the Adriatic cultures was further investigated by imputing the genotypes in 815 (new and published) samples [8–10,18–26] and generating a PCA based on the shared identity-by-descent (IBD) fragments between individuals (Additional file 2: Table S10; Additional file 3: Fig. S7A). Notably, we confirmed a significant shift of the Adriatic people toward the Balkan and Central European populations with respect to Etruscans and Italy_IA_Romans (Additional file 3: Fig. S7B).

### Social structure and mobility in pre-Roman Central Italy

By performing genetic kinship analyses, we did not identify close relatives (1st and 2nd degree) among our samples, not even within the well-represented Novilara necropolis (Additional file 2: Table S2 and S3). This result may reflect the fact that Novilara was probably one of the largest Picene funerary sites [11,12] and possibly served a huge inhabited area where it is more difficult to identify relatives.

To get a finer insight into the social structure of ancient Italian groups, we used their imputed genomes to extract the Runs of Homozygosity (ROHs) longer than 4 cM (Additional file 2: Table S11). While the highest levels of ROHs are found within Mesolithic individuals (mean of ROHs = 206.57 cM), later groups show lower levels of inbreeding (mean of ROHs = 24.12 cM), as previously highlighted [18,27] (Additional file 2: Table S10; Additional file 3: Fig. S8). Notably, although ROH distribution of the Picenes is comparable to other IA groups (mean of ROHs in Picenes = 42.85 cM; Etruscans = 17.23 cM; Italy_IA_Romans = 47.63 cM; Italy_IA_Apulia = 23.65 cM, ANOVA *p*-value = 0.164), there are two individuals (PN125 and PN24) with a F_RoH_ value (0.111 and 0.096, respectively), suggesting inbreeding events in their genealogical history (Additional file 2: Table S12).

Among the central Italic IA populations analyzed here, we observed the presence of several genetic outliers (i.e., individuals deviating from their main PCA/Admixture cluster; Additional file 3: Fig. S1 and S9). In Novilara, PN103 and especially PN172 are partially shifted towards Near Eastern populations in the PCA and show a high proportion of the CHG/Iran Neolithic component in Admixture analysis (Fig. 2B). For PN172, the Y chromosome reflects this finding, since it belongs to the Near Eastern haplogroup J1-M267/Z2223 (Additional file 2: Table S13; Additional file 3: Fig. S10). On the other hand, PN50 seems to be more shifted towards Caucasian and/or Central European populations (Additional file 3: Fig. S1 and S9).

Despite the absence of a clear pattern, outgroup f3-statistics analysis in the form f3(X, Y; YRI.SG) (where X is an outlier individual and Y a modern or ancient Eurasian population) shows that PN103 and PN172 are more similar to European Neolithic populations (Additional file 2: Table S14; Additional file 3: Fig. S11). Notably, in the qpWave/qpAdm framework, PN103 and PN172 have a higher proportion of (and a higher number of feasible models with) Near Eastern and North African ancestral components (i.e., Levant_PPN or Morocco_EN) with respect to the general Picene population (Additional file 2: Table S8). On the contrary, PN50 shows great affinity with Central and Northern European IA populations with respect to other Novilara individuals, possibly indicating genetic influences from these regions (Additional file 3: Fig. S11). Moreover, qpWave/qpAdm highlights a greater Yamnaya influence in this individual; indeed, when it is modeled as a mixture between Serbia_IronGates_Mesolithic, Italy_N and Yamnaya, the last component is up to about 50% higher compared to the same model for the general Picene population (Additional file 2: Table S8).

Other Novilara individuals (PN13, PN20, PN43 and PN137) and one Sirolo-Numana sample (PSD5) are partially separated from the main Picene cluster, greatly overlapping with the variability of Etruscans (Additional file 3: Fig. S1 and S9). However, with outgroup f3-statistics and qpWave/qpAdm analyses, it is not possible to highlight major differences compared to the general Picene cluster (Additional file 2: Table S8 and S14; Additional file 3: Fig. S11).

In the Etruscan site of Monteriggioni/Colle di Val D’Elsa, we found an outlier (EV15A) that is more similar to modern Sardinians or European Neolithic populations and a sample (EV18) showing a greater North European ancestry, similarly to other Etruscan outliers previously reported [9] (Fig. 2; Additional file 3: Fig. S1 and S9). In outgroup f3-statistics, EV15A seems to be rooted in the genetic variability of pre-Bronze Age Europe, being very similar to populations that lack (or have a small proportion) of the Yamnaya-like component (Additional file 2: Table S14; Additional file 3: Fig. S11). Whereas EV18 is similar to populations with a high proportion of this component, like the Czech Iron Age individuals (Additional file 2: Table S14; Additional file 3: Fig. S11). These observations are mirrored in qpWave/qpAdm analysis; in the model with Serbia_IronGates_Mesolithic, Italy_N and Yamnaya components, the last one is found with a slight lower proportion in EV15A with respect to the general Etruscan population (23% vs 25%), while in EV18 it is extremely high (53%) (Additional file 2: Table S8).

### The genetic legacy of the Roman Empire in the Middle Adriatic area

To study the evolution of the gene pool after the Romanization of the former Picene territory we analyzed 7 individuals of the Late Antiquity Pesaro site (less than 6 km from Novilara) (Fig. 1; Additional file 2: Table S1). In the PCA we observe a substantial shift towards modern and ancient Near Eastern populations in the genetic landscape of Adriatic Central Italy as already reported for other parts of the peninsula since the Roman Imperial period (Fig. 2A; Additional file 3: Fig. S1 and S7) [8,9,15]. We confirmed these results also with the Admixture analysis (Fig. 3B), where a great genetic influence from the CHG/Iran Neolithic component is observed in almost all Pesaro individuals.

Nevertheless, some differences among individuals can be observed. Indeed, two of them, namely PF1 and PF32, are strongly shifted towards North Africa and Near East in the PCA, respectively (Additional file 3: Fig. S1 and S9). Consistently, in the Admixture analysis they show no signs of the EHG/Yamnaya component but have a high proportion of the CHG/Iran Neolithic one (Fig. 2B). On the contrary, PF15 has no CHG/Iran Neolithic ancestry, while it shows a very high proportion of the EHG/Yamnaya component possibly as a consequence of genetic continuity with previous IA groups or a more recent Central/Northern European ancestry (Fig. 2B).

While the outgroup f3-statistics did not indicate a clear pattern (Additional file 3: Fig. S11), these results were supported by the qpWave/qpAdm framework (Additional file 2: Table S8). For the general population of Pesaro, several models show ancient Near Eastern genetic components (e.g., CHG, Iran Neolithic, Levant PPN) as the most represented ancestries. When performing the same models on the outliers, PF1 and PF32 have a greater amount of Near Eastern ancestry, while PF15 has usually less with respect to the general population. Notably, while some models including Morocco_EN can explain the ancestry of both the general Pesaro population and the outliers, only in the case of PF1 this component reaches a non-negligible proportion (≈10%), suggesting a possible past genetic influence from North Africa for this individual (Additional file 2: Table S8).

The great genetic influence from the Near East in Pesaro (and in general in post-IA Italian populations) is mirrored also in IBD (Additional file 3: Fig. S7) and f4/NNLS analysis in which all the Italian Imperial/Late Antique sites here analyzed cluster together (Additional file 3: Fig. S6).

### Phenotypic shifts in Italy from the Copper Age to modern times

We investigated possible shifts in phenotypic traits through different ages and places by imputing 111 markers related to pigmentation, metabolism and immune response in both the ancient specimens analyzed in this study and ancient samples available from the literature, for a total of 874 individuals from the Copper Age (CA) to the Medieval period (Additional file 2: Table S15, S16 and S17). The allele frequency shifts in the phenotypic markers have been evaluated by comparing: 1) all the Italian groups from the CA to the modern period, (Additional file 2: Table S16), and 2) all the populations dated to the 1^st^ millennium BCE from all the geographic areas here considered (Additional file 2: Table S17).

Interpreting these results with due caution because of the general small sample size, we observed 7 markers showing significant allele frequency differences in both tests. Three of these SNPs (namely, rs3135388 in the *HLA* locus, rs2395182 on *HLA-DRA* and rs2066842 in *NOD2*) are involved in the immunity response. Focusing on the two *HLA* markers, when comparing all the 1^st^ millennium BCE groups (test 2), the significance is mainly driven by the Italy_IA_Romans, showing the lowest frequency of the effective alleles (Additional file 2: Table S16). Interestingly, when comparing all the Italian populations (test 1), we observed a high frequency of both *HLA* effective alleles since the CA and in all the IA groups except Italy_IA_Romans. Later, we observe a decrease in their allele frequencies, suggesting a homogenizing effect of Roman domination in the Italian peninsula for these loci, which later show a new allele frequency increase after the Medieval period.

As for the third SNP rs2066842, for which the effective allele is considered a risk factor for Chron’s disease [28], its significance is due to the Picene group that shows the highest frequency of the effective allele compared to all the Italian populations over time and to the other coeval ones.

The other 4 significant variants in both tests are in the *SLC45A2* and *HERC2* loci and are involved in pigmentation, with their allele frequency changes showing a temporal and geographic pattern in line with the prediction of darker eye, hair and skin colors (Additional file 2: Table S15, S16 and S17). Interestingly, the Picenes have a greater proportion of individuals with blue eyes (30.2%) and blond hair color (20.9%) than other Italic populations. In the Etruscans and Italy_IA_Romans these lighter phenotypes are much less common (blue eyes: 2.6% in the Etruscans, 10.0% in the Italy_IA_Romans; blonde or dark blond hair: 5.3% in the Etruscans, 10.0% in the Italy_IA_Romans), making these populations more similar to previous individuals from the Italian peninsula.

In the statistical test involving all the Italian groups (test 1), we also observed 10 significant differences in the allelic frequency in SNPs involved in carbohydrate and vitamin metabolism, immunity and pigmentation. In particular, two of these SNPs are in the *MCM6/LCT* locus and have been associated with lactase persistence in adulthood in Europe [29]. These markers show an overall low frequency of the lactase persistence allele in Italy through time, increasing only from the Late Antiquity and Medieval period, possibly due to the additional foreign genetic inputs, as shown by the PCA and Admixture analysis. In any case, with few exceptions, possibly caused by random fluctuations, Picene people are overall similar to the other Iron Age Italian populations.

## Discussion

In this study we analyzed the genetic variation among 54 Central Italian samples spanning more than 1,000 years of history and portrayed a comprehensive picture of the evolution of the genetic pool of the pre-Roman middle Adriatic cultures by contextualizing them in the broader framework of the entire Italic Iron Age, providing novel insights into the population dynamics after the Roman Empire in the area.

### The Picenes in the context of Central Italian IA

We present the first genetic characterization of the Picene ethnicity, highlighting a substantial homogeneity among the sites analyzed here (Novilara and Sirolo-Numana). Our genetic data align with archaeological evidence suggesting that the Picenes emerged as an ethnic group from earlier Italic Bronze Age cultures (Fig 2A, Additional file 3: Fig S1). This finding is particularly interesting because, despite sharing a core material culture, some differences in the archaeological record of the two Picene sites here analyzed can be observed [5]. Our results show that these differences were only cultural, as the whole Picene genetic pool exhibits either a common origin or a significant genetic homogenization (Fig. 2). However, we caution that given the small sample size from Sirolo-Numana, further investigations involving more individuals from different necropolises are needed. More in general, despite the high cultural diversity within the Italian context [1,4], our analysis has revealed strong genetic homogeneity in the Iron Age ethnic groups suggesting that their genetic origins are interconnected (Fig. 2A, Additional file 3: Fig. S3).

One of the most interesting characteristics of the Picenes and of all the Italic IA populations analyzed so far is the frequent presence of genetic outliers (Fig. 4). In Novilara two individuals (PN172 and PN103) show a greater Near Eastern ancestry compared to the general population. These individuals may represent a direct movement of people from the Eastern Mediterranean area to the Middle Adriatic region, as also attested from an archaeological perspective which indicates clear evidence of the circulation of goods and cultural patterns [13]. Another Novilara individual (PN50) is more compatible with a Central European ancestry, possibly representing genetic connections associated with the extensive trade network between Picenes and Central-Northern Europe where personal ornaments with a hint of Picene influence have been discovered [13]. This finding suggests that, within Novilara society, there were individuals with different origins who were well-integrated into it. This is further supported by the archaeological analysis of the grave goods in the outliers’ tombs, which shows no differences from the broader Picene culture. Similarly, two genetic outliers (EV15A and EV18) can be observed among the Etruscans from Monteriggioni/Colle Val D’Elsa. While the former shows a genetic makeup analogous to coeval Sardinians [22], probably representing the genetic outcome of the well-known IA connections between this island and Etruria [30], the latter is more similar to Central European BA/IA individuals, like other Etruscan outliers already described [9]. The presence of all these genetic outliers in the Italic IA, also attested among the Romans [8], is a direct consequence of the great mobility of the Mediterranean populations [7,31]. This trend started from the Neolithic and intensified throughout the subsequent periods, reaching considerable levels in the 1^st^ millennium BCE [32,33] and with the Italian peninsula, due to its central position in the Mediterranean basin possibly representing a melting pot for these movements. Thus, the increased mobility in the IA resulted in a highly multicultural society which profoundly affected the Italian cultures [1,3–5]. Our results highlighted the importance to analyze several samples from the same cultural background, in order to unveil the many social aspects characterizing them. Moreover, our findings confirm that a process of globalization was already well in place before the Roman Imperial times [8].

**Figure 4.**
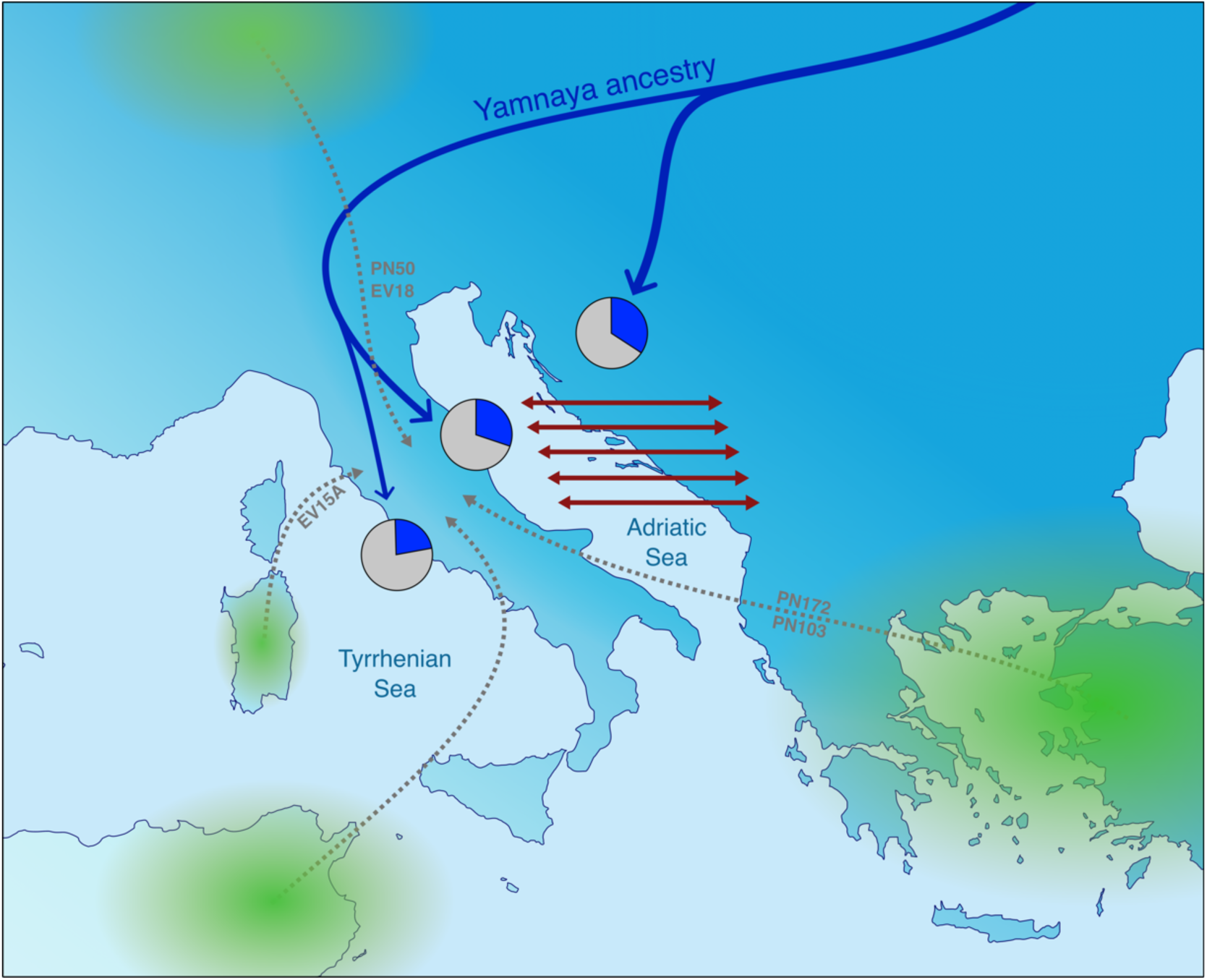
Proposed scenarios for the genetic evolution of the Italic IA. The differences observed between Italic IA Adriatic and Tyrrhenian populations, mainly represented by different proportions in the Yamnaya-related ancestry (blue color in the pie charts) could be explained by two scenarios: 1) differential arrival of the Yamanaya-related ancestry (blue arrows); 2) trans-Adriatic genetic connection represented (red arrows). Genetic outliers observed in the Central Italic IA (this study, Antonio et al. (2019), Posth et al. (2021)) and their putative area of origin (based on genetic ancestry, green gradients) are depicted.

It is worth noting the absence of relatives among Picene individuals here analyzed, in particular in the Novilara necropolis. This can be explained considering that this necropolis is one of the largest Picene funerary sites excavated so far [7,12]. This indicates that, if present, the corresponding inhabited area may have had a considerable extension. Nevertheless, in Novilara we identified at least two individuals (PN24 and PN125) with an amount of ROHs consistent with the hypotheses that they are offspring of two related individuals, possibly first cousins (Additional file 2: Table S12; Additional file 3: Fig. S8). Therefore, even though close relatives were not directly identified in this study, social and kinship relationships may have been complex and far from being completely resolved.

### Corridors and barriers: The Adriatic Sea and the Apennines

As highlighted in several studies, the amount of Yamnaya-related genetic component varies greatly in Europe starting from the BA, with Southern European populations (i.e., Italian and Balkan peninsulas) usually showing less of it. [18,34–39]. In this context, it is worth noting that our results showed some genetic similarity between Italy and Balkans, with the IA Italy and BA/IA Balkans being sister clades (Additional file 2: Fig. S6). This relative genetic homogeneity between these peninsulas (Additional file 3; Fig. S1) probably started much earlier than the BA according to previous studies [36,40] and seems to have been maintained during the BA and IA. Here, we propose two non-mutually exclusive scenarios for the genetic similarity between Italian and Balkan populations (Fig. 4): 1) These areas were influenced by similar demographic events, mostly involving the arrival of the Yamnaya-related genetic component from continental Europe along the two sides of the Adriatic Sea; 2) The two peninsulas were in close genetic contact during the BA and the IA. The populations on the Adriatic Sea could have moved between the two shores and mixed with each other, in a continuous process of gene flow.

These scenarios can also explain the small differences observed between the Adriatic and Tyrrhenian Italic IA populations determined by the East to West genetic gradient encompassing the Italian and Balkan peninsulas, mostly represented by the Yamnaya-related genetic ancestry (Additional file 3: Fig. S5). Indeed, the arrival of the Yamnaya component in Italy, possibly through its North-Eastern regions, may have been partially hindered by Apennine mountains, resulting in a higher Yamnaya genetic component in Adriatic populations compared to Tyrrhenian ones. In addition, and considering the second scenario, if gene flow occurred across the Adriatic Sea during the BA and/or IA, it would certainly have impacted more the populations facing the sea, and then gradually diminished as distances from it increased. Future studies focusing on the genetic onset of the BA on both Adriatic shores (Italian and Balkan) compared to the Tyrrhenian side may help to elucidate these processes.

From an archaeological perspective, the extensive connections across the two peninsulas throughout the BA and IA are well-characterized. Strong commercial trans-Adriatic routes were already present from the 3^rd^ millennium BCE [7]. During the Early BA the Cetina culture, although rooted in the Dalmatian coast, spread throughout the Adriatic, eventually reaching Sicily, Malta and Western Greece [7,41,42]. These contacts persisted throughout the BA [43] and during the IA they were strongly consolidated. Indeed, the extensive presence of shared cultural traits across the two sides of the Adriatic Sea has allowed some authors to describe an “Adriatic *koiné*” (Adriatic culture) to emphasize this circulation of goods and perhaps individuals [44–46]. Similarly, the possible genetic relationship between Northern/Central Europe and the Middle Adriatic region could be supported by the observed material connections between the Hallstatt culture along the Danube River and Northern-Central Italy, already starting from the Late BA [47].

Y chromosome data of the Italic IA groups provide additional evidence to these observations, suggesting that the two scenarios proposed are complementary. Indeed, in the Picenes, two main Y haplogroups are observed, namely R1-M269/L23 (58% of the total) and J2-M172/M12 (25% of the total) (Additional file 1: Table S13), which may be representative of the direct connection to Central Europe and the Balkan peninsula, respectively. As for the R1-M269/L23 haplogroup, it has been associated with the Yamnaya ancestry [19] and observed at high frequency among Central European populations from the BA onward. More specifically, one Picene individual (PN13) clusters with modern and ancient Central-Northern Europeans and other IA Italians, in the sub-branch defined by the L51/L11 markers, frequent in mainland Europe [48] (Additional file 2: Fig. S10). Another Picene individual (PN62) belongs to the R1-L23/Z2106 subclade, which has been previously interpreted as a genetic link between Yamnaya, Balkans and Southern Caucasus [19]. Finally, five Picenes and two Etruscans are placed at the basal portion of the R1-L23 branch, together with other ancient Yamnaya, Balkan and Southern Caucasic samples (Additional file 2: Fig. S10). On the other hand, it is worth noting that the trans-Adriatic distribution of the internal branches of J2-M172/M12 was previously interpreted as a clue of a BA expansion from the Balkans in the Italian area and a link between BA Balkans and BA Nuragic Sardinia, possibly with peninsular Italian intermediates that were not observed before [19,49]. Interestingly, two out of three of our J2-M12 Picene samples (PN91 and PN101), due to their phylogenetic position (Additional file 3: Fig. S10) in between the BA Nuragic and the BA Balkan clusters, could represent the descendants of the aforementioned Italian intermediates.

Overall, combining genetic and archaeological evidence, we suggest that the Adriatic Sea was a hotspot for the commercial, social and genetic connections between the two peninsulas, with the populations living on its shores directly involved in the exchanges. It is possible that in the highly connected landscape of the Mediterranean IA, mountain ranges like the Apennines may have been a greater barrier to the movement of people than small stretches of sea like the Adriatic one. Nevertheless, it is possible that other factors (e.g., cultural barriers) contributed to the emerging picture of the Italic IA genetic pool.

### After the Romanization

With the onset of Late Antiquity, here represented by the Pesaro necropolis, a substantial shift in the genetic landscape of the former Picene area towards the Near East can be observed. This process is the genetic outcome of the social changes brought by the Romanization of the Italian peninsula in general. Indeed, in Italy, Near Eastern genetic influx, represented by the CHG/Iran Neolithic component, starts to be extensively present in the majority of the Imperial time individuals, and it continues to be common during the Late Antiquity [8,9], as observed also for Pesaro necropolis in this study (Fig. 2B). The most likely explanation for this genetic shift is the central role of Rome and Italy in the political and social scene of the Roman Empire, attracting people from the other Imperial provinces, especially the wealthy Eastern ones [8].

Footprints of the initial spread of the Romans across the peninsula and its impact on demographic changes can be observed in the frequency shift of some alleles associated with phenotypic traits. Indeed, the allele frequency at two *HLA* markers associated with protection against leprosy and risk factor for gluten intolerance [50,51] were probably influenced by the Roman expansion. They are found with high frequency in all the Italian groups from the Neolithic to the IA, apart from the Roman IA individuals. In subsequent periods, the frequencies drop for all the Italian areas considered (Additional file 1: Table S16), suggesting a strong homogenizing effect caused by the Roman domination. Only during the Middle Ages, the frequency of these alleles rose again, suggesting an intricate evolutionary history for these markers, intertwined between demographic dynamics and putative selective pressure which possibly had a role in the diffusion of gluten intolerance as a consequence of increased protection against leprosy.

Despite the diffusion of Near Eastern ancestries seems to be relatively homogeneous and widespread in the Italian context, in Pesaro necropolis we observe a great genetic variability, towards both the extremes of the distribution [8,9]. This may be the result of population dynamics occurring in the Mid-Adriatic area during Late Antiquity. After the fall of the Western Roman Empire, the area was under control of the Eastern Roman (Byzantine) Empire during the 6^th^ century CE, with the city of Pesaro representing an important political center of the Duchy of the Pentapolis. It is possible that these political dynamics additionally influenced the movement of people in the area and, therefore, the evolution of the genetic pool.

It was only in later periods, starting from the Early Middle Age, that the Italian genetic pool changed again, with a decrease of Near Eastern ancestry and a new increase of Central/Northern European one. The main reason for this is probably the massive arrival of people with a Central European ancestry (like the Longobards) that established the nowadays North-South genetic gradient in Italy [8,52,53].

## Conclusions

Our study provides new insights into the population dynamics in Italy from the Iron Age to the Late Antiquity. In particular, we investigated demographic events occurring in the Adriatic side of Central Italy in more than 1,000 years of history and throughout several socio-political changes. We identified a common genetic origin for all the Italian IA ethnicities analyzed until now that can be traced back to the arrival of the Yamnaya-related ancestry starting from the Bronze Age. We highlighted the genetic similarities of the Italian and Balkan peninsulas during these ages, indicating common histories and/or frequent contacts across the Adriatic. Our results show the presence of several genetic outliers among the Picenes, as it has also been identified in other IA groups, suggesting that a cosmopolitan society began to emerge and persisted in Italy during the IA, reaching its climax during the Roman Imperial period. With the onset of the Roman Empire in the area and in the subsequent Late Antiquity, we observed a shift in the genetic landscape toward the Near East mirroring the pattern observed in Rome and on the Tyrrhenian side, pointing out that this change likely affected the entire peninsula.

## Methods

DNA extraction and library preparation were performed at the ancient DNA laboratory at the Estonian Biocenter, Institute of Genomics, University of Tartu, Tartu, Estonia. Quantification and sequencing of the libraries were carried out at the Estonian Biocenter Core Laboratory.

### DNA Extraction

We extracted DNA from a total of 102 human remains samples, consisting of 3 petrous bones and 99 tooth roots. Small slices of bone were sampled from petrous bone and root portions were taken from teeth samples. Both these procedures were performed with a sterile drill wheel that was sterilized with 6% (w/v) bleach followed by distilled water and ethanol rinse in between the samples. The collected samples were placed in 6% (w/v) bleach for 5 minutes, then rinsed with 18.2 MΩcm water for 3 times and soaked for 2 minutes in 70% (v/v) ethanol. During the previous bleach and ethanol steps the samples were shaken to allow the detachment of particles. Later, the samples were placed on a clean paper towel inside a class IIB hood, and they were left to dry for two hours with the UV light on. To calculate the correct volume of EDTA and Proteinase K needed for the extraction, the samples were weighted. We considered 20x EDTA [µL] of sample mass [mg] and 0.5x Proteinase K [µL] of sample mass [mg]. The samples, EDTA and Proteinase K were placed into PCR-clean conical tubes (Eppendorf) of 5 or 15 mL, depending on the total volume required, under the IIB hood. These tubes were incubated on a slow shaker for 72 h at room temperature. The resulting DNA extracts were concentrated to a final volume of 250 µL with the Vivaspin Turbo 15 (Sartorius). Then, they were purified in large volume columns using the High Pure Viral Nucleic Acid Large Volume Kit (Roche) with 2.5 mL of PB buffer, 1 mL of PE buffer and 100 µL of EB buffer (MinElute PCR Purification Kit, QIAGEN). The silica columns were placed in a collection tube to dry and later in a 1.5 mL DNA lo-bind tube (Eppendorf) for elution. Samples were incubated at 37 °C for 10 minutes with 100 µL of EB buffer and then centrifuged at 13,000 rpm for 2 minutes. The silica columns were removed after centrifugation and the samples were stored at −20 °C, 30 µL of the samples were used for library preparation.

### Library preparation and sequencing

The libraries for sequencing were made with NEBNext DNA library Prep Master Mix Set for 454 (E6070, New England Biolabs) and with Illumina-specific adaptors [54] using established protocols [54–56]. The end repair part was implemented (as described in Saupe et al. [18]) using 118.75 µL of water, 7.5 µL of buffer and 3.75 µL of enzyme mix, incubated at 20°C for 30 minutes. Then, the samples were purified with 500 µL of PB buffer and 650 µL of PE buffer and eluted in 30 µL of EB buffer (MinElute PCR Purification Kit, QIAGEN). As previously, the adaptor ligation step was implemented as in Saupe et al. [18], using 10 µL of buffer, 5 µL of T4 ligase and 5 µL of adaptor mix [54], incubating for 14 minutes at 20°C. The samples were purified as described above, and then eluted in 30 µL of EB buffer (MinElute PCR Purification Kit, QIAGEN). The step of the adapter fill-in was performed using 13 µL of water, 5 µL of buffer and 2 µL of Bst DNA polymerase, incubating 30 minutes at 37°C and for 20 minutes at 80°C [18]. For PCR amplification of the libraries, we used 50 µL of DNA library, 1X PCR buffer, 2.5 mM MgCL2, 1 mg/mL BSA, 0.2 µM inPE1.0, 0.2 mM dNTP each, 0.1 U/µL HGS Taq Diamond and 0.2 µM indexing primer. Cycling conditions were settled in the following way: 5 s at 94°C, followed by 18 cycles of 30 s each at 94°C, 60°C and 68°C, with a final extension of 7 minutes at 72°C. After PCR amplification, the samples were purified in 35 µL of EB buffer (MinElute PCR Purification Kit, QIAGEN). To measure the concentration of dsDNA/sequencing libraries and to confirm that library preparation was successful, we performed three verification steps: fluorometric quantification (Qubit, Thermo Fisher Scientific), parallel capillary electrophoresis (Fragment Analyzer, Agilent Technologies) and qPCR. DNA sequencing was performed using the Illumina NextSeq500/550 High-Output single-end 75 cycle kit and 20 samples were sequenced together on one flow cell.

### Mapping

The sequences of the adapters, the indexes, the poly-G tails that occur because of the NextSeq 500 technology specifics and sequences shorter than 28 bp (--minimum-length 28, to reduce the risk of random mapping of sequences from other species) were removed from the DNA sequences before the mapping with cutadapt-2.1 [57]. The resulting sequences were mapped to the human reference sequence GRCh37 (hs37d5) with BWA-0.7.17 [58]. To reduce the effect of reference bias in the following analyses we used the command bwa aln with relaxed alignment parameters (-n 0.01 -o 2) in combination with disabling seeding (-l 1024) [59–61]. The sequences were converted to BAM file format subsequent to the alignment and only the mapped sequences were retained using samtools-1.9 [62]. Duplicates were removed with picard-2.20.8 (http://broadinstitute.github.io/picard/index.html) and indels were realigned using GATK-3.5. With samtools-1.9 we filtered out reads with mapping quality lower than 25 as suggested in Martiniano et al. [59]. We estimated the number of final reads, average read length, average coverage and other parameters with samtools-1.9 on the final BAM files. The endogenous DNA content of the samples (calculated as the proportion reads mapping to the human reference genome) spanned between 0,16% to 69,34% with an average of 22,33% (Additional file 2: Table S1).

### aDNA authentication and contamination rate

We employed the program MapDamage-2.0 [63] to estimate the frequency 5’ ends of sequences C->T transitions, one of the characteristic patterns of ancient DNA (aDNA) damage, to confirm that the sequences we obtained are mostly ancient. We estimated contamination rates on the mitochondrial DNA (mtDNA) with the method detailed in Jones et al. [64] by computing the fraction of non-consensus bases at mtDNA haplogroup defining position. Samples with a contamination rate lower than the 3% were used for subsequent analyses.

### Genetic sex estimation

Genetic sex was estimated with the procedure described in Skoglund et al. [65], computing the proportion of reads mapping to the Y chromosome with respect to the total number of reads mapping either to the X or the Y chromosome. In some cases, it was not possible to unequivocally assign the genetic sex, but we note that these samples were not considered for population genomics analysis because of other issues (i.e., low coverage or contamination, Additional file 2: Table S1).

### mtDNA haplogroups assignment

We assigned the mitochondrial DNA haplogroups with the online tool Haplogrep2 [66] (https://haplogrep.i-med.ac.at/haplogrep2/index.html). The VCF file that was used to perform haplogroup prediction was obtained calling the variants from the .bam files with bcftools-1.14 [67]. The command used was bcftools mpileup with the additional flag --ignore-RG and then only the variant positions were called with the command bcftools call -m --ploidy 1 -v.

### Y chromosome haplogroup assignment and phylogeny

Y chromosome phylogenetic relationship among newly reported samples and other ancient Euroasiatic samples [8–10,18,19,21–26,68–70] (Additional file 2: Table S5) were reconstructed with pathPhynder [71] starting from the .bam files, using standard parameters. As a reference tree, we used the one provided in Martiniano et al. [71] which span across all the genetic variability of human Y chromosome haplogroups with a total of 2014 individuals included. To visualize the resulting tree, we used the R package ggtree [72].

### Variant calling on autosomes

Variant calling for autosomes was performed with ANGSD-0.917 [73]. We called haploid genotypes sampling a random base (-doHaploCall 1) for each position that is present in the 1240K SNPs panel using the -sites options. In addition, we specified the major and minor alleles as they are indicated in the 1240K SNPs panel with the option -doMajorMinor 3. The function haploToPlink was used to convert the .haplo file, resulting after the SNP calling, into Plink [74] format files.

### Design of datasets for genomic analysis

For genomic analyses, we compared our newly reported samples with modern and ancient individuals from the literature. The included samples belong to the AADR dataset v52.2 ([16], https://reich.hms.harvard.edu/allen-ancient-dna-resource-aadr-downloadable-genotypes-present-day-and-ancient-dna-data) both the 1240K and the 1240K+HO dataset, Lazaridis et al. [19] and Aneli et al. [10]. To maximize the number of SNPs available for each analysis we built three different dataset: A) including 1240K and HO data, used for PCA and Admixture analysis; B) including only 1240K data, used for kinship analysis, f3-statistic, D-statistics, qpAdm; C) including 1240K, HO and modern Italians from Raveane et al. [52], employed for PCA to compare ancient and modern Italians (Additional file 3: Fig. S2). All the data coming from different datasets were merged with Plink-1.9 [74]. To minimize the possible error due to post-mortem damage of ancient DNA, for population genetics analysis we kept only transversions from these datasets. The resulting total number of autosomal SNPs was: 98,845 for dataset (A), 209,089 for dataset (B) and 98,842 for dataset (C). After merging, we retained for subsequent analysis only the newly reported samples showing mtDNA contamination lower than 3% and a coverage of SNPs of at least 10,000 from dataset (A) and 5,000 from dataset (B) and (C). The label for the population assigned to each individual sample used in this study can be found In Additional file 1: Table S4 and S5. Because they belong to the same ethnicity and for the genetic similarities outlined, for several analysis (D-statistic, qpWave/qpAdm, f4/NNLS, IBDs, ROHs and phenotype prediction) Etruscan individuals reported in this study were grouped together with individuals from the literature [9], maintaining outliers individuals separated.

### Kinship analysis

Kinship estimations were performed independently with READ [75] and TKGWV2 [76]. Since READ with default parameters estimates the median pairwise distance of unrelated individuals in the population under study to assess the potential kinship relationships, using different individuals may change these estimates. Therefore, for READ we performed tests with four different datasets: 1) samples from central Italy Iron Age [8,9], but excluding the ones identified as outliers; 2) only the Picenes, with both the necropolises of Novilara and Sirolo-Numana; 3) only the Picenes from the necropolises of Novilara; 4) the LA individuals from Pesaro with the Imperial time individuals from Antonio et al. [8]. For each of these tests we performed the analysis with all the SNPs and with only the transversions (Additional file 2: Table S2). The samples were selected from the original datasets with the option --keep of Plink [74] and converted into the .tped format that is required for READ. With TKGWV2 we performed two different tests with all SNPs and only transversions including all the samples of interest at the same time (newly reported individuals, other IA samples from central Italy and Imperial time individuals), since it requires an external source for allele frequencies estimation. We used the allele frequencies of the European populations of the 1000 Genomes data [77] as provided here: https://github.com/danimfernandes/tkgwv2 (Additional file 2: Table S3). In all the tests performed the two Etruscan samples EV7A and EV19 resulted to be the same individuals (or monozygotic twins), therefore for population genetics analysis the sample with the lowest coverage (EV19) was excluded.

### Principal component analysis

For PCA we selected a subset of modern and ancient Eurasian and North African samples from dataset A, for a total of 1464 individuals (Additional file2: Table S4). The subset was performed with the option --keep of Plink [74]. The resulting Plink format files were converted into EIGENSTRAT format with the program convertf of the EIGENSOFT-7.2.0 package [78,79] using the parameter “familynames:NO”. The program smartpca of the EIGENSOFT-7.2.0 package was used to perform PCA with the parameters lsqproject:YES, autoshrink:YES and outliermode:2. We projected all the ancient individuals onto the PCs built based on modern samples. In the PCA performed including modern Italians (employing dataset C, Additional file 3: Fig. S2), these had to be projected like ancient individuals due to the high number of missing SNPs (>60%), since the original data were produced on a different beadchip with respect to the HO and 1240K data (Infinium Omni2.5-8 Illumina beadchip). The results of the first two PCs were visualized in R-4.1.3 (https://www.r-project.org/) with the package ggplot2 (https://ggplot2.tidyverse.org/).

### Admixture analysis

We performed unsupervised Admixture analysis [17] with 1708 modern and ancient Eurasian and North African individuals selected from dataset A (Additional file 2: Table S4). Before running Admixture, SNPs were pruned for linkage disequilibrium with Plink [74] (option --indep-pairwise with parameters: 50 5 0.5) for a final set of 98759 transversions. Moreover, modern and high-coverage ancient diploid genomes were converted into pseudo-haploid by picking a random allele for each variant position. We performed 10 independent repetitions (giving different random seed with the -s option) for k values from 2 to 10 using the option --haploid=’*’. The results from different runs were merged and visualized with R package pophelper (Additional file 3: Figure S3) [80]. We portrayed k=4 because it is the most representative, showing a differentiation between CHG and Yamnaya components.

The interpolation maps with the proportion of the ancestral components obtained with Admixture were performed with QGIS-3.26.1 (https://www.qgis.org/en/site/). For this analysis we kept only the samples from the I millennium BCE excluding known outliers for a total of 396 samples and, in archaeological sites where more than one individual was present, we performed the mean for each component (Additional file 2: Table S7). The interpolation was calculated with the IDW method and for representation the option “singleband pseudocolor” was chosen, the minimum value was set to 0.01 and the maximum to 0.6 and, finally, the option “Interpolation: Discrete” was selected.

### Outgroup f3 statistic

To explore the genetic relationships between different IA Italian groups, the Pesaro LA individuals and the putative outliers identified with the PCA and/or Admixture analysis in the form f3(X,Y;YRI.SG), where X is one of the following population: Picene, Etruscan, Italy_IA_Romans, Italy_IA_Apulia, Pesaro_LA, Italy_Imperial; or one of the following samples: EV15A, EV18, PN103, PN13, PN137, PN172, PN20, PN43, PN50, PF1, PF15, PF32; Y is a modern or ancient Eurasian or North African population selected from the dataset B (Additional file2: Table S4). f3 statistics were computed with the program qp3pop of the package admixtools-7.0.1 [81] using the option “inbreed:YES”.

### D-statistic

D-statistic was performed in the form D(X, Y; Italy_BA_EBA, YRI.SG), where X and Y are a set of the populations present in Additional file 2: Table S4, selected from the dataset B. An additional test comparing Italy_IA_Tyrrhenian and Italy_IA_Adriatic populations was added. D-statistic was calculated with the program qpDstat of the package admixtools-7.0.1. [81] with the options “printsd: YES” and “inbreed: YES”.

### qpWave/qpAdm analysis

To explore more in detail the ancestral genetic components of the Italian IA and LA populations, and the putative outliers we identified, we exploited a qpWave/qpAdm framework [82]. As a target, we used the populations and single individuals in the following list: Picene, Etruscan, Italy_IA_Romans, Italy_IA_Apulia, Pesaro_LA, Italy_Imperial, EV15A, EV18, PN103, PN13, PN137, PN172, PN20, PN43, PN50, PF1, PF15, PF32. In detail:

i) we exploited qpWave to evaluate if the targets can be described as a combination of the sources.
ii) if (i) returned a *p*-value > 0.01, we used qpAdm to model the targets as a mixture of the sources.

We tested models with two and three left populations (sources) using all the possible combinations of the following: Anatolia_N_Barcin, Serbia_IronGates_Mesolithic, Yamnaya, Morocco_EN, Levant_PPN, CHG, Iran_N, Italy_N, Sardinia_N, Germany_BellBeaker, Italy_CA, Italy_BA_EBA, EHG. We discussed plausible models with a *p*-value ≥ 0.01 as also reported in Skourtanioti et al. [83]. As right populations (outgroup) we always used the following: Mbuti.DG, ISR_Natufian_EpiP, Morocco_Iberomaurusian, Mesopotamia, Russia_AfontovaGora3, Russia_MA1_HG.SG, Turkey_Boncuklu_N, Turkey_Epipaleolithic, WHG2, Ethiopia_4500BP.SG (the corresponding samples can be found in Additional file 2: Table. S4).

### f4 NNLS analysis

To explore at a finer level the ancestral components of the newly reported samples and other populations from the literature, we performed a Non-Negative Least Squares (NNLS) analysis exploiting different f4-statistics vectors as a proxy for the relationships among the analyzed populations. In detail, we performed approximately 1.5 millions of f4 in the form f4(X,Y;Z,Mbuti.DG) where X, Y and Z are all the possible combinations of 112 populations. For each of these populations, a f4 vector was created and used to perform NNLS analysis (reconstructing each target individual copying vector as a mixture of different proportions of the putative sources employing a slightly modified version of the nnls function in the R package ‘‘nnls’’, as described in Saupe et al. [18] and Wangkumhang et al. [84]) with these two sets of ancestral populations:

1) Anatolia_N_Barcin, Serbia_IronGates_Mesolithic, Iran_N, Levant_PPN, Yamnaya;
2) Anatolia_N_Barcin, Serbia_IronGates_Mesolithic, Iran_N, Levant_PPN, EHG.

The issue with this method is that some NNLS values are not computed from f4 vectors, in particular when the same populations are present between the X, Y and Z populations. To avoid the presence of missing data, we set all these f4 values to 0, which is formally correct only when X and Y, and not X and Z, are the same populations. Additionally, to get an overview of the genetic similarity among the analyzed populations we performed an UPGMA tree based on the Euclidean distance between f4 values.

### Genome imputation

We performed genome imputation following a slightly different pipeline to the one described in Hui et al. [85]. In particular, genotypes were called with ANGSD-0.917 [73] on the SNPs panel (removing indels) present in the global population of the 1000 Genomes Project Phase 3 [77], using the parameters -doMajorMinor 3 -GL 1 -doPost 1 -doVcf 1 -doMaf 1 -checkBamHeaders 0. Genotype likelihoods were updated in Beagle-4.1 [86] with -gl mode. Then, we performed imputation from sites where the genotype probability (GP) of the most likely genotype is equal or higher than 0.99, using Beagle-5 [87] with -gt mode. We exploited the 1000 Genomes Project Phase 3 [77] world population as a reference for the Beagle -gl step, while we used the Human Reference Consortium [88] dataset for the Beagle -gt step. After genome imputation, an additional GP filter (MAX(GP) ≥ 0.99) was applied before performing subsequent analyses.

We selected for imputation only samples with a coverage ≥ 0.1X, for shotgun data, or covering more than 300K SNPs from the 1240K panel, for chip data. To compare our data with other ancient populations, we selected unrelated individuals from different studies [8–10,18–26] for a total of 815 individuals. The complete list of samples and the corresponding publication can be found in Additional file 2: Table S5).

### Identity-by-descent (IBD) analysis

In order to identify segments of the genome that are identical by descent, we exploited the program IBDseq-vr1206 [89] on the imputed dataset, following the procedure illustrated in Ariano et al. [90]. In detail, with Plink-1.9 [74] we filtered out SNPs showing genotype missingness > 0.02 and a MAF < 0.05 (parameters --geno 0.02 and --maf 0.05 in Plink). The resulting Plink format files were converted into VCF with the option --vcf of Plink, and this was used as the input file for IBDseq. The parameters used for IBDseq were errormax = 0.005 and LOD >= 3, as indicated in Ariano et al. [90] and Schroeder et al. [91], and only IBD longer than 2Mb (≈2cM) were retained as suggested in Browning and Browning [89]. Individuals included in IBD analysis are listed in Additional file 2: Table S5. A PCA was performed with the data of IBD sharing between each pair of individuals in the dataset with the function prcomp and the parameters center=TRUE, scale.=TRUE in R. Results were visualized with R package ggplot2.

### Runs of homozygosity

To identify possible traces of past inbreeding events in ancient Italian samples, we performed runs of homozygosity analysis using hapROH [92]. In order to run hapROH with the reference data provided in Ringbauer et al. [92], we downsampled the dataset obtained after genome imputation to the autosomal SNPs in the 1240K SNPs panel, with Plink-1.9 [74] option --extract. Data were converted into EIGENSTRAT format with the program convertf of the EIGENSOFT-7.2.0 package [78] using the parameter “familynames:NO”. hapROH analysis was performed with parameters e_model="haploid" and random_allele=True. Results were visualized with R package ggplot2. The complete list of samples on which we performed ROH analysis can be found in Additional file 2: Table S5.

### Phenotype prediction

In order to perform phenotype prediction, we ran a two-step pipeline for local imputation of the region around the phenotypic markers of interest, as previously described [85]. In details, for 39 out of the 41 HIrisPlex-S set of SNPs [93], we selected 2 Mb around the informative variants, merging the regions on the same chromosome, except for the variants on chromosome 15, which have been analyzed in two different regions since the distance between the two nearest SNPs was about 20 Mb. We selected 10 regions from 9 autosomes, spanning from about 1.5 Mb to 6 Mb. For the other phenotypic informative markers (diet, immunity and diseases), we selected 2 Mb around each variant and merged the overlapping region, for a total of 46 regions from 17 autosomes. We performed this phenotypic analysis on the same set of samples used for the whole genome imputation described above.

We called the variants using ANGSD-v0.917 [73] at positions with a minimum allele frequency (MAF) ≥ 0.1% in the reference panel, that was composed by the Europeans from the 1000 Genomes (EUR) [77] plus the MANOLIS (EUR-MNL) set from Greece and Crete extracted from the HRC [88]. The ANGSD output in VCF format was used as input for the first step of our imputation pipeline [85] (genotype likelihood update), performed with Beagle 4.1 -gl command [86] using the same panels as before as reference. We then discarded the variants with a genotype probability (GP) lower than 0.99 and imputed the missing genotype with the -gt command of Beagle 5.1 [87] using the HRC as a reference panel. We then discarded the variants with a GP < 0.99 and used the remaining SNPs to perform the phenotype prediction. For the pigmentation prediction, we prepared an input file for the HIrisPlex-S webtool (https://hirisplex.erasmusmc.nl) following its manual for formatting and results interpretation. Sample-by-sample phenotype prediction and genotypes (as counts of the effective alleles in the form 0, 1 or 2) are reported in Additional file 1: Table S15. We then grouped the individuals in different cohorts depending on both time and space. First, we grouped the ancient individuals from the Italian peninsula in 10 groups from the CA to the Medieval period and compared them with the modern TSI from the 1000 Genomes Project. We compared the groups performing an ANOVA test and, for the significant variants, we performed a *post-hoc* Tukey test to identify the significantly different pairs of groups (Additional file2: Table S16). Using the same approach, we also analyzed the difference among populations dated to the 1^st^ millennium BCE, i.e., coeval to the Picene people, creating 15 groups (Additional file2: Table S17). For both comparisons, we set a significance threshold applying a Bonferroni’s correction on an alpha value of 0.05 divided by the number of tested SNPs.

## Availability of data

The dataset generated and analyzed during the current study are available in the European Nucleotide Archive (ENA, https://www.ebi.ac.uk/ena/browser/home) under Accession Number PRJEB62031.

## Abbreviations

IA: Iron Age
BA: Bronze Age
CA: Copper Age
BCE: Before Common Era
CE: Common Era
IBD: Identity-by-descent
ROH: Runs of Homozygosity

## Ethics approval and consent to participate

Not applicable

## Consent for publication

Not applicable

## Authors’ contributions

FRa, ED, CD and BT conceived the study; CD, SF, PG and EC collected the samples and provided archaeological background; FRa, HN and AS performed aDNA extraction and sequencing; FRa, LDG, FM, RH, MH, LP, Fri, CG and ED performed bioinformatic analyses; CLS, KT, MM, ED, FC and BT supervised the work; FRa, FM, CLS, KT, ED, FC and BT discussed and interpreted the results; FRa, ED, FC and BT acquired the funding; FRa, ED and BT wrote the original manuscript with input from all the authors. All authors read and approved the final manuscript.

## Acknowledgements

Bioinformatic analyses were carried out employing the facilities of the High Performance Computing Center of the University of Tartu. The authors are grateful to the aDNA team at the Institute of Genomics of the University of Tartu, Estonia, including Lehti Saag and Biancamaria Bonucci for the support in wet lab and bioinformatic analysis. The authors are grateful to Superintendence Archaeology, Fine Arts and Landscape of the Marche Region and to Superintendence Archaeology, Fine Arts and Landscape of the Toscana region. LDG was supported by #NEXTGENERATIONEU (NGEU) and the Ministry of University and Research (MUR), National Recovery and Resilience Plan (NRRP), project MNESYS (PE0000006, DN. 1553 11.10.2022). FM was supported by Fondazione con il Sud (2018-PDR-01136) and by MUR (2022P2ZESR). The EASI-Genomics project has received funding from the European Union’s Horizon 2020 research and innovation programme under grant agreement No 824110.

