## Additional file 1 for "The Genomic portrait of the Picene culture: new insights into the Italic Iron Age and the legacy of the Roman expansion in Central Italy"

**Additional file 1: Information about the necropolises analyzed**

***Novilara necropolis***

The Novilara necropolis (province of Pesaro and Urbino) [1,2] is located on the northern slope of a small hill at about 4 km from the Adriatic coast. The Novilara site, one of the largest and best-documented Picene funerary contexts [3], provides outstanding evidence of an Iron Age necropolis and it has been known in literature since the late 19^th^ century. Between 1892 and 1893, Edoardo Brizio discovered 263 burials dating back to the 8^th^ and 7^th^ centuries BCE [3,4].

In 2012 new research began, as a consequence of some major works related to the enlargement of a motorway. The area under investigation (10.000 sq. meters) corresponds to the “Molaroni area” of Brizio’s excavations. During the 2012-2013 excavations, 150 new burials have been brought to light, dating back to the 8^th^ and 7^th^ centuries BCE. These new graves are the subject of this study and are characterized by a variable amount and type of items including vessels, weapons, pins, necklaces, and amber earrings, as well as tools related to weaving and spinning (e.g., spools, spindle whorls, spindles).

Except for three male cremations, placed inside urns only partially preserved, the graves preserve single inhumates laid on their sides. There are also three double burials of persons deposited at the same time and one double burial with inhumates buried at different times [1]. Traces of shrouds covering the whole body were found in most burials, while wooden coffins surrounding the bodies were observed only in some inhumations, although it is still not possible to identify a specific pattern associated, for example, with a specific social status [1]. There are sectors of the necropolis that are more densely used and others where the burials are arranged in irregular alignments or in small semicircular groups, separated by gaps of different widths, which could also indicate the presence of possible paths within the necropolis [1]. Some of the burial clusters are so well defined and separated from the other tombs to suggest the presence of family, clan, or other kind of non-biological strong relationships among the inhumated. Several grave goods made of bronze, iron and ceramics were found in different burials, in some cases indicating established connections with other cultural groups. Some burials are richer than others, indicating a non-egalitarian society and thus the possible emergence of an elite. This is particularly evidenced by the presence of child burials with rich grave goods, clearly indicating that their wealth was inherited from the high social class to which they belonged and not from a status acquired during their lifetime [1,2].

***Sirolo-Numana necropolis***

The funerary area of Sirolo-Numana is located very close to the modern city of Ancona. Several burials, covering a time span of the entire Iron Age, have been found in this area, located in different points of the territory and far apart from each other [1]. Since the late 7^th^ century BCE, in this area it is possible to observe burial clusters that were posed inside big circular ditches [1]. These clusters appear in the archaeological record following great social changes in the Picene society. They probably represent the emergence of a war-rooted aristocracy, manifesting its prestige with large family tombs [1]. Similar to Novilara, in this area the bodies were deposed inside a shroud, and in some cases also in a wooden coffin, above a layer of gravel probably of marine origin [1]. Although this burial ritual is found in other Picene necropolises, it has been present at this site since the earliest periods of Picene civilization and is not correlated with specific gender, age, or social status [1]. It seems that some kind of markers were indicating the burials, although the many visible overlaps suggest that older graves were respected and not violated only for a limited period of time. However, it could also indicate the willingness to continuously use the same burial area to emphasize the bonds to a specific group [1].

***Monteriggioni/Colle di Val d’Elsa necropolis***

We generated data from the Etruscan necropolis of Monteriggioni/Colle di Val d’Elsa. This necropolis, also known as “Necropoli del Casone” is located between the Tuscan towns of Monteriggioni and Colle di Val d’Elsa. This necropolis was used as a burial site from the Iron Age to the Roman period (8^th^ century BCE-2^nd^ century CE), as testified by the different layers and type of tombs. The samples analyzed in this study come from one of the latest discovered burials in this necropolis, a chamber tomb dated to the “Orientalizing phase” (8^th^-6^th^ century BCE) of the Etruscan culture, characterized by a heavy influence from the eastern Mediterranean and Near Eastern cultures. According to the luxury furniture, jewels, objects, and decoration found in this chamber, it has been suggested that it was an aristocratic burial.

***Pesaro necropolis***

Pesaro necropolis is located in present-day via Cavallotti, Pesaro, Marche region [5]. The area represented one of the suburban areas of the ancient city of *Pisaurum*, placed 200 meters south of the city walls, close to the ancient Roman road *Via Flaminia*. The Late Antiquity part of the necropolis here analyzed consists of a total of 39 inhumations. Most of the tombs do not show grave goods and they are found in different structure types. In particular, “*alla cappuccina*” tombs, box tombs in brick, and wood lined earthen graves can be observed. It is possible to observe two different groups of tombs, considering topographical and stratigraphic data. The main group is composed of burials oriented in a northwest-southeast direction, arranged in approximately regular rows and all built with “*alla cappuccina*” or tile box structures. The second group is composed of the pit graves located in the northern part of the site and of the “*alla cappuccina*” tombs with substructure in masonry. These burials are oriented in a partially different direction compared to the first group. Moreover, they cross the road (*Via Flaminia*), disrupting some of the tombs from earlier periods and with the fill of the drainage trenches. In some of the burials in the earthen pits, it was present a simple grave good, i.e. a non-adorned two-toothed comb made in bone placed on the skull, the shoulder, the chest, or the hand. In addition, a pair of bronze earrings and a necklace with beads in bone and in glass paste were found in a burial attributed to a woman, dating this tomb to a time spanning from the 6^th^ and 7^th^ century CE. Except for these objects, it was not possible to retrieve any additional dating element. Thus, chronological attribution was based on stratigraphy. The initial date is determined by the presence of ceramic material in the drainage trenches at both sides of the road which was cut through by the more recent tombs. Ceramics that were common between the end of the 6th century and the beginning of the 7^th^ century (in particular, fragments of Middle Adriatic *terra sigillata*, cooking pots (*ollae*) with engraved decorations, fragments of rimmed bowls, a few fragments of Oriental amphorae, and late-production stoppers) identify a narrow time range between the renovation of the road and the subsequent development of the burials. Therefore, these findings may indicate that the first burial group was realized during the 6^th^ century CE, when the road was still in use, while the second group can be probably dated to the early 7^th^ century CE, when the road was no longer used.

For its location and dating, Pesaro necropolis can be placed in the context of Byzantine Italy, being part of the Duchy of the Pentapolis, one of the regions in which the Exarchate of Ravenna, a newly established province of the Eastern Roman Empire, was subdivided. It was called Duchy of the Pentapolis because its power was spread across two series of five cities: one on the sea (comprising the cities of Pesaro, Rimini, Ancona, Senigallia and Fano); and one on the mountain side (comprising the cities of Gubbio, Cagli, Urbino, Fossombrone and Jesi). This Byzantine province was conquered by the Longobards between 727 and 729 CE. Pope Stephen II, in order to weaken Longobards power in Central Italy and strengthen his political influence in the region, sought the help of Pepin the Short, the first Carolingian king of Franks. The Frank military campaign in Italy was successful, and the territory belonging to the former Duchy of the Pentapolis was donated to the pope, contributing to the emergence of the Papal State [6].
