## Additional file 3 for "The Genomic portrait of the Picene culture: new insights into the Italic Iron Age and the legacy of the Roman expansion in Central Italy"

### Additional file 3: Supplementary figures

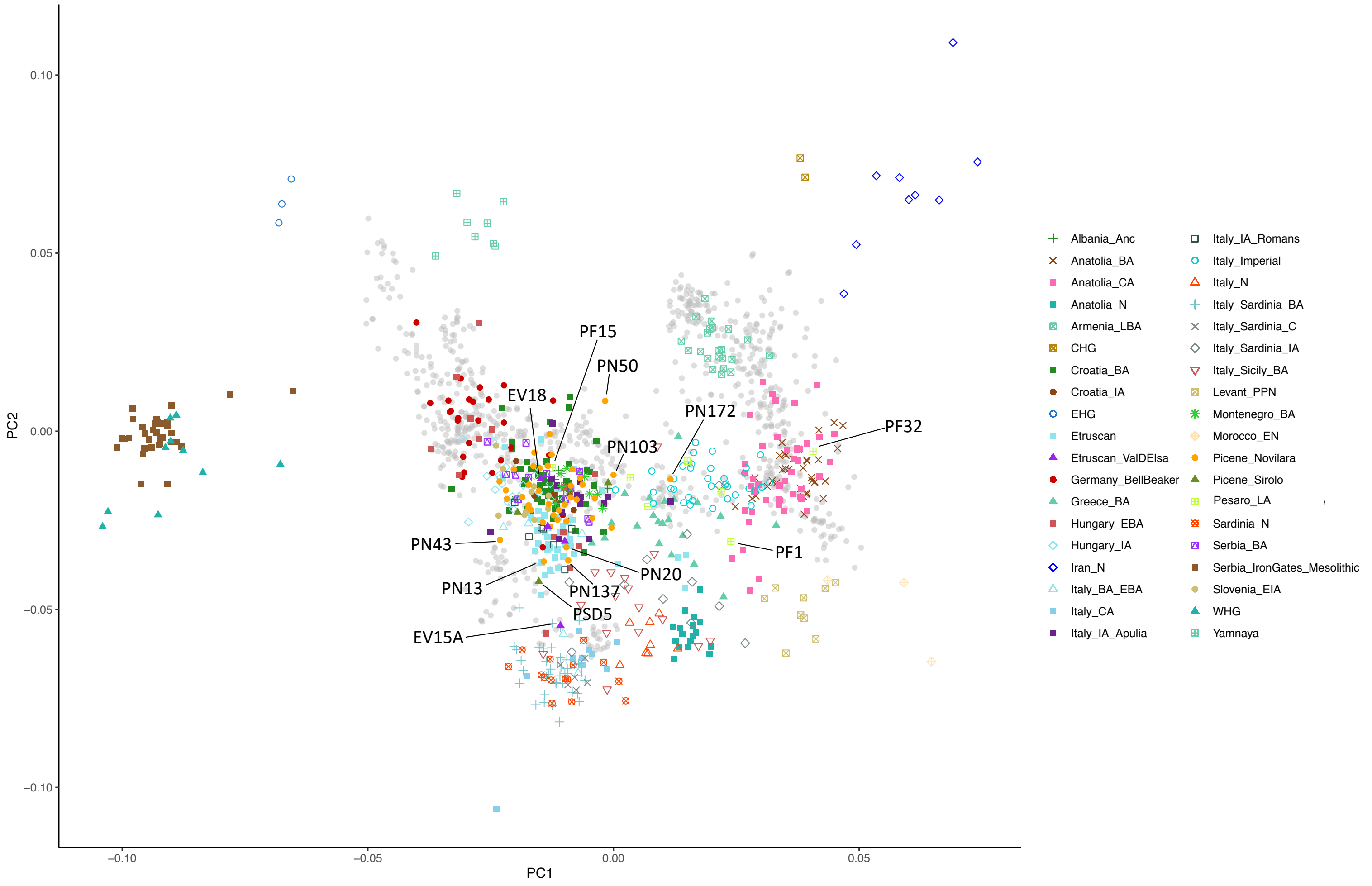


**Figure S1.** Complete version of the PCA shown in figure 2 of the main text. The putative genetic outliers are highlighted. Modern individuals are in gray and not shown in the legend.

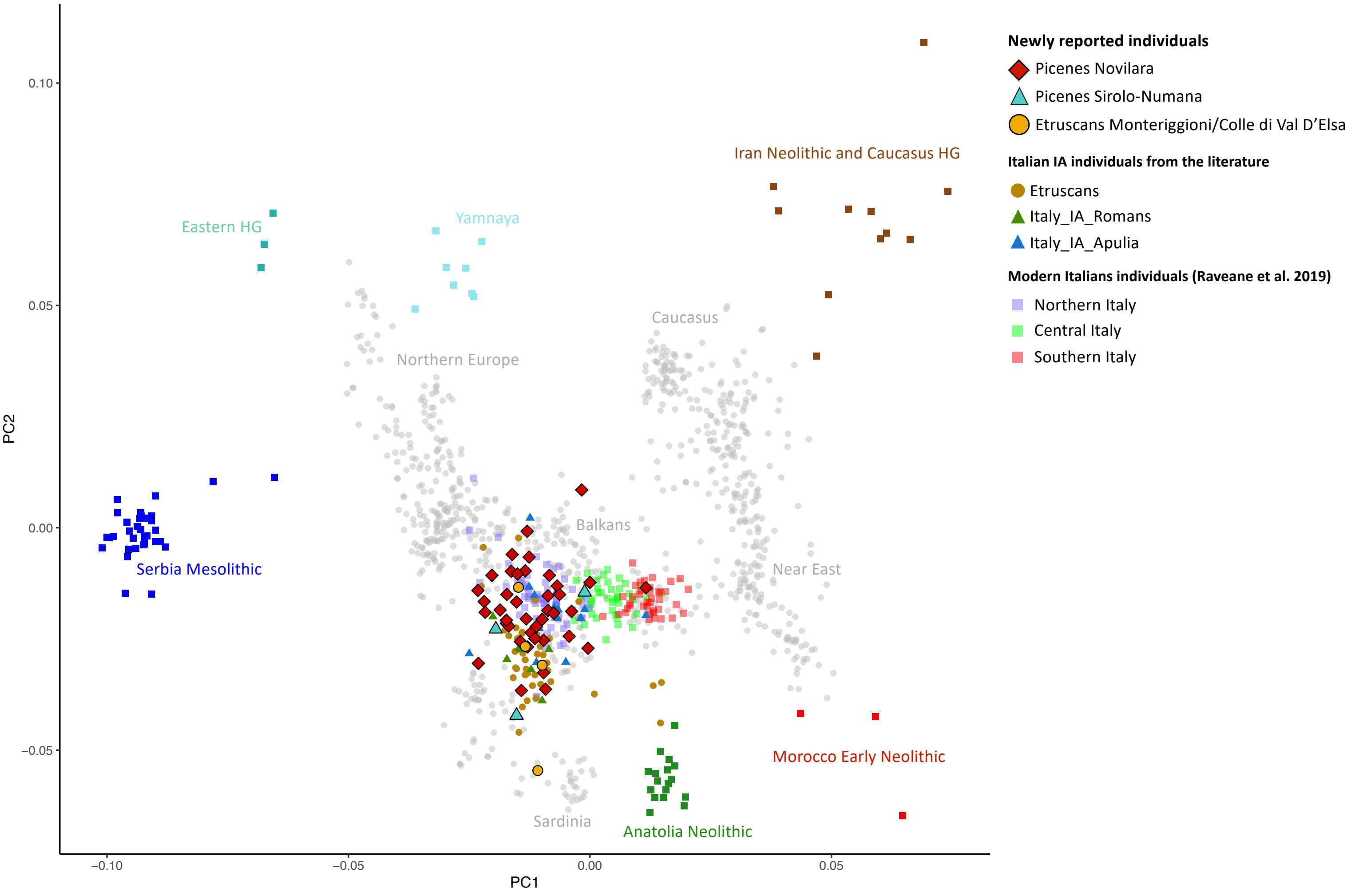


**Figure S2.** PCA including Italic IA individuals and modern peninsular Italians from northern, central and southern regions. Modern Italians data from Raveane et al. (2019).

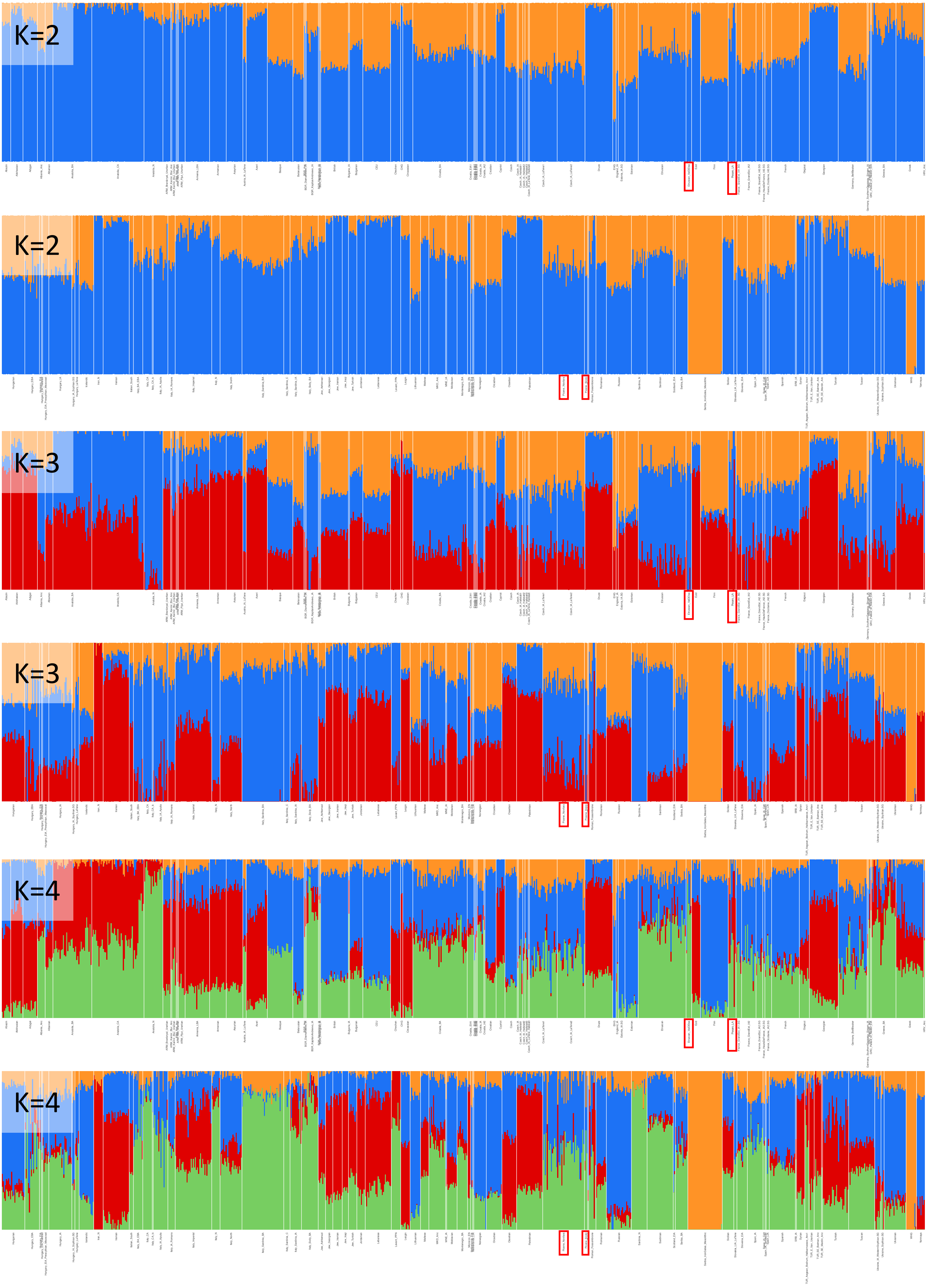


**Figure S3.** Unsupervised Admixture analysis from k=2 to k=4 with all the 1708 modern and ancient samples included in the analysis. Newly reported individuals are highlighted by red squares.

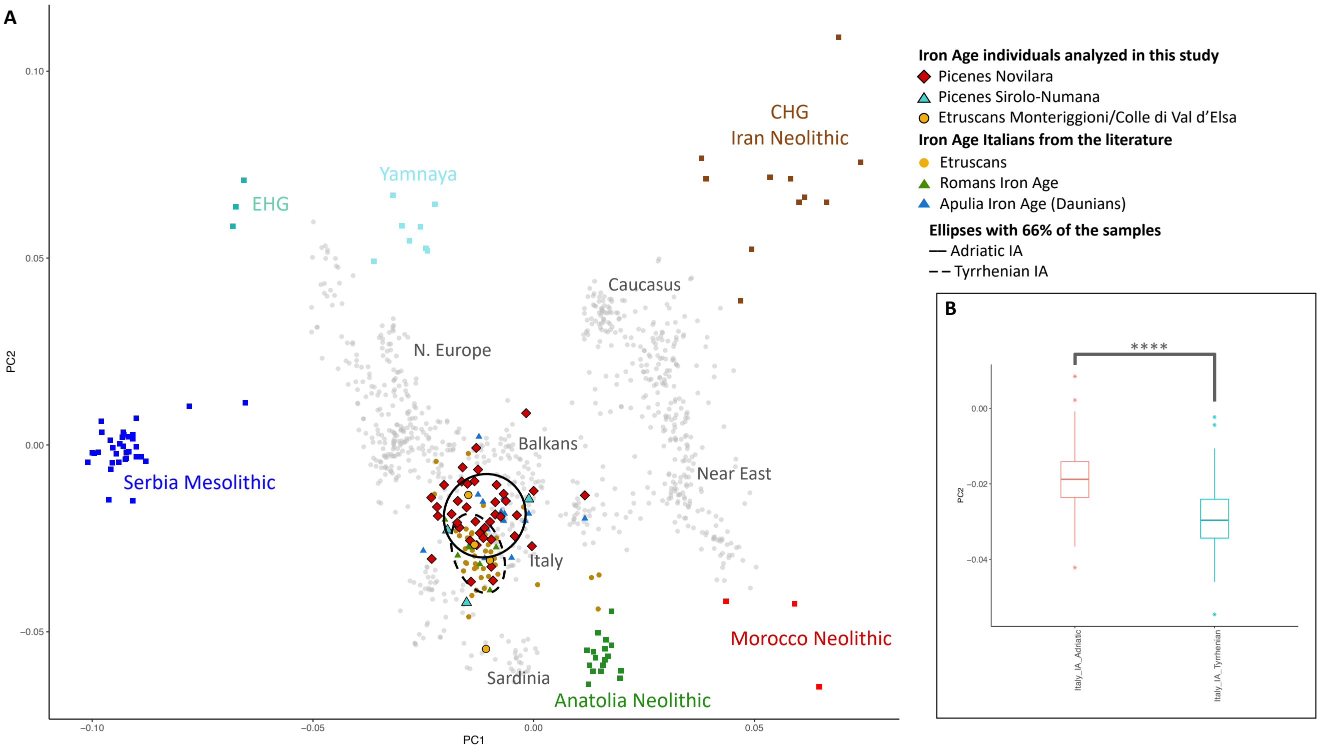


**Figure S4.** A) Tyrrhenian and Adriatic differences in the PCA in figure 2 of the main text. Ellipses drawn considering 66% confidence level for a multivariate t-distribution of the Adriatic and Tyrrhenian IA individuals are represented, highlighting the shift of the Adriatic populations towards Yamnaya individuals. B) Boxplots representing the distribution along the PC2 of the Italic IA individuals from the Adriatic and the Tyrrhenian side. Colored asterisk in the plot indicate outliers. Significance was calculated with the Mann-Whitney U Test: p-value < 0.0001.

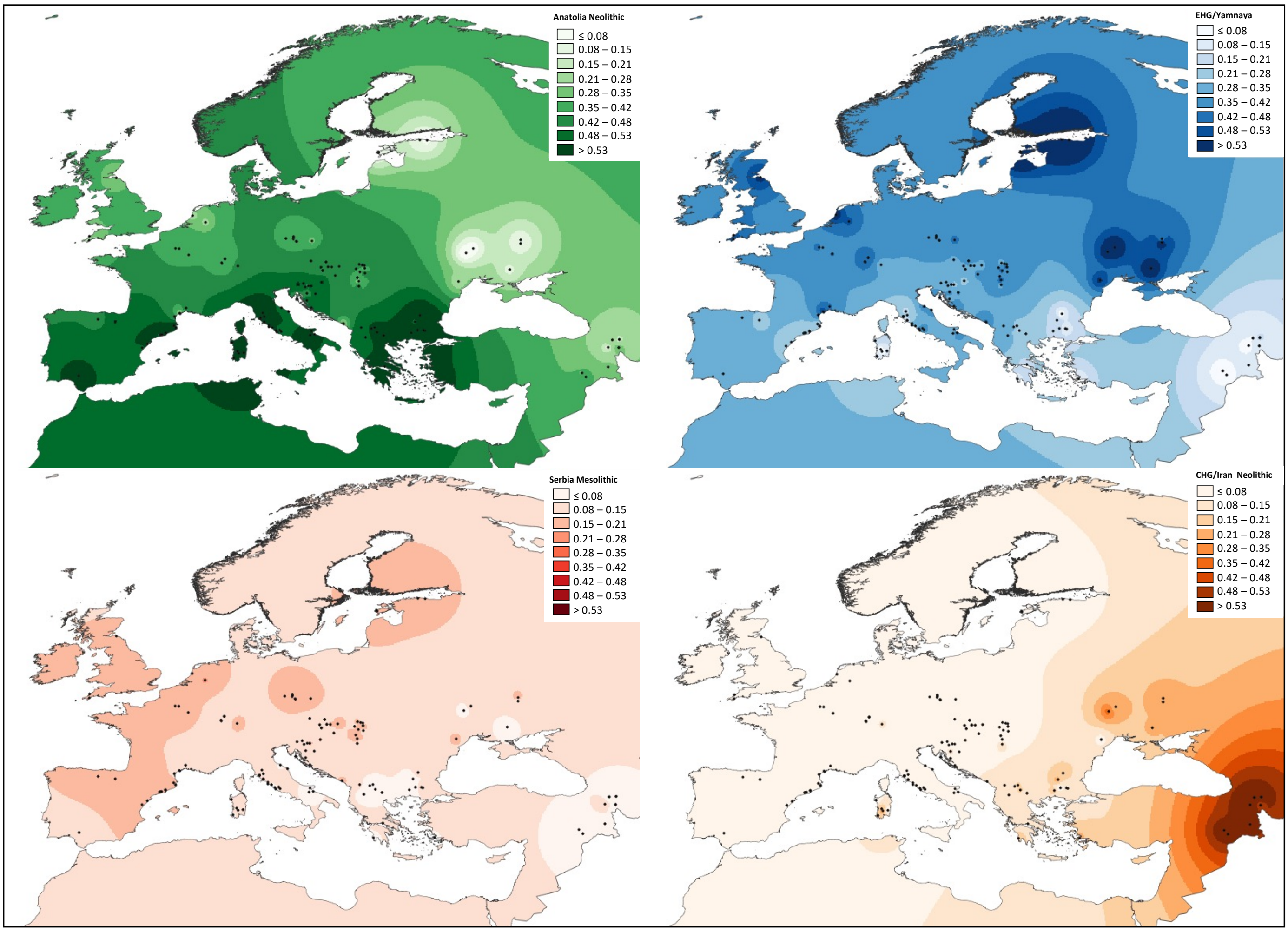


**Figure S5.** Interpolation maps of the ancestral Eurasian components. Black dots: 1st millennium BCE samples (from present study and from the literature, for details see Additional file 2: Table S7). When more samples were present in a site, the average of the proportion of each component was computed. Strong outlier individuals were excluded from this analysis.

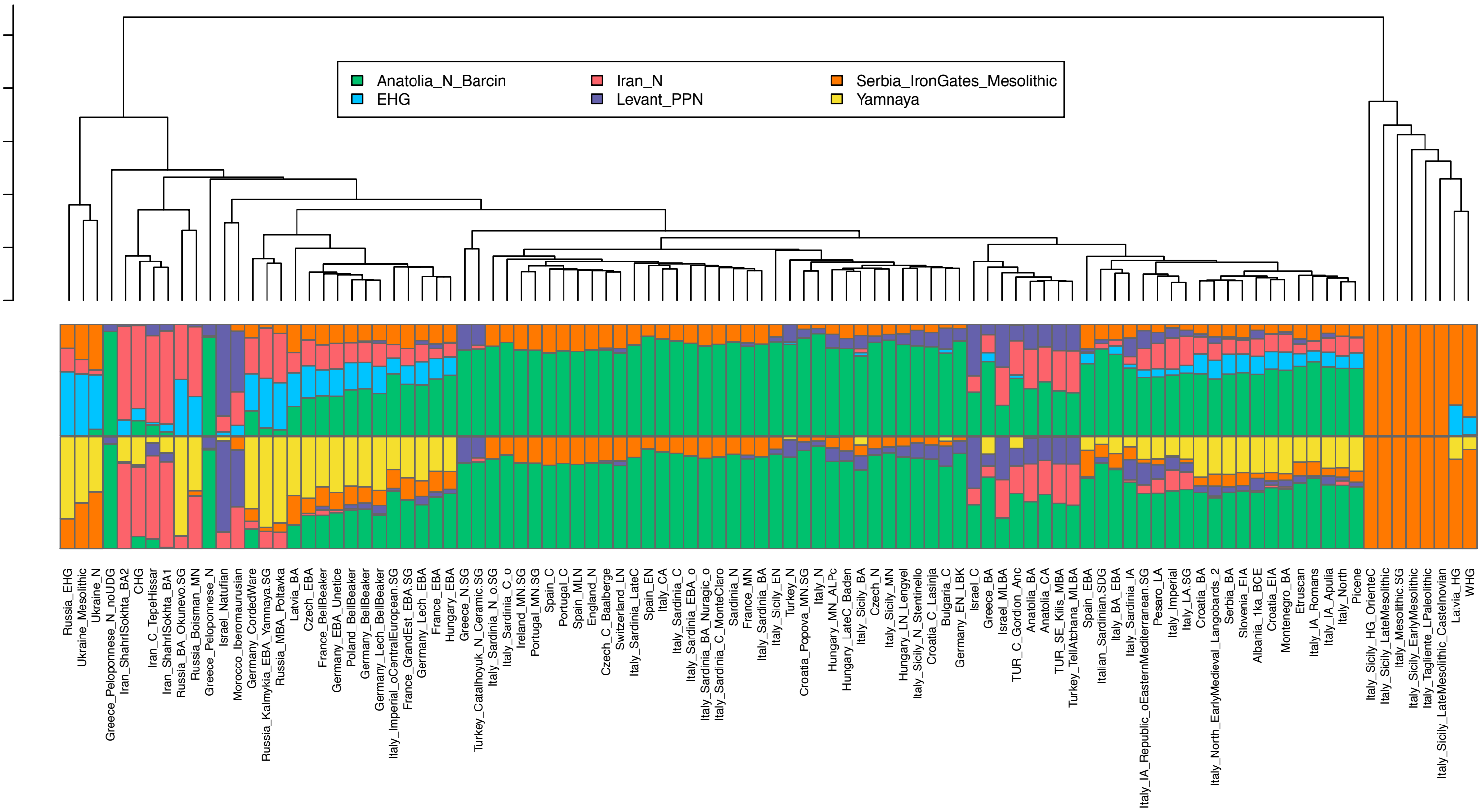


**Figure S6.** f4/NNLS analysis with two sets of ancestral genetic components, one including the EHG ancestry (first bar plot) and the other the Yamnaya one (second bar blot). On the top, the UPGMA tree highlights the genetic similarities between the populations analyzed based on the f4/NNLS vectors.

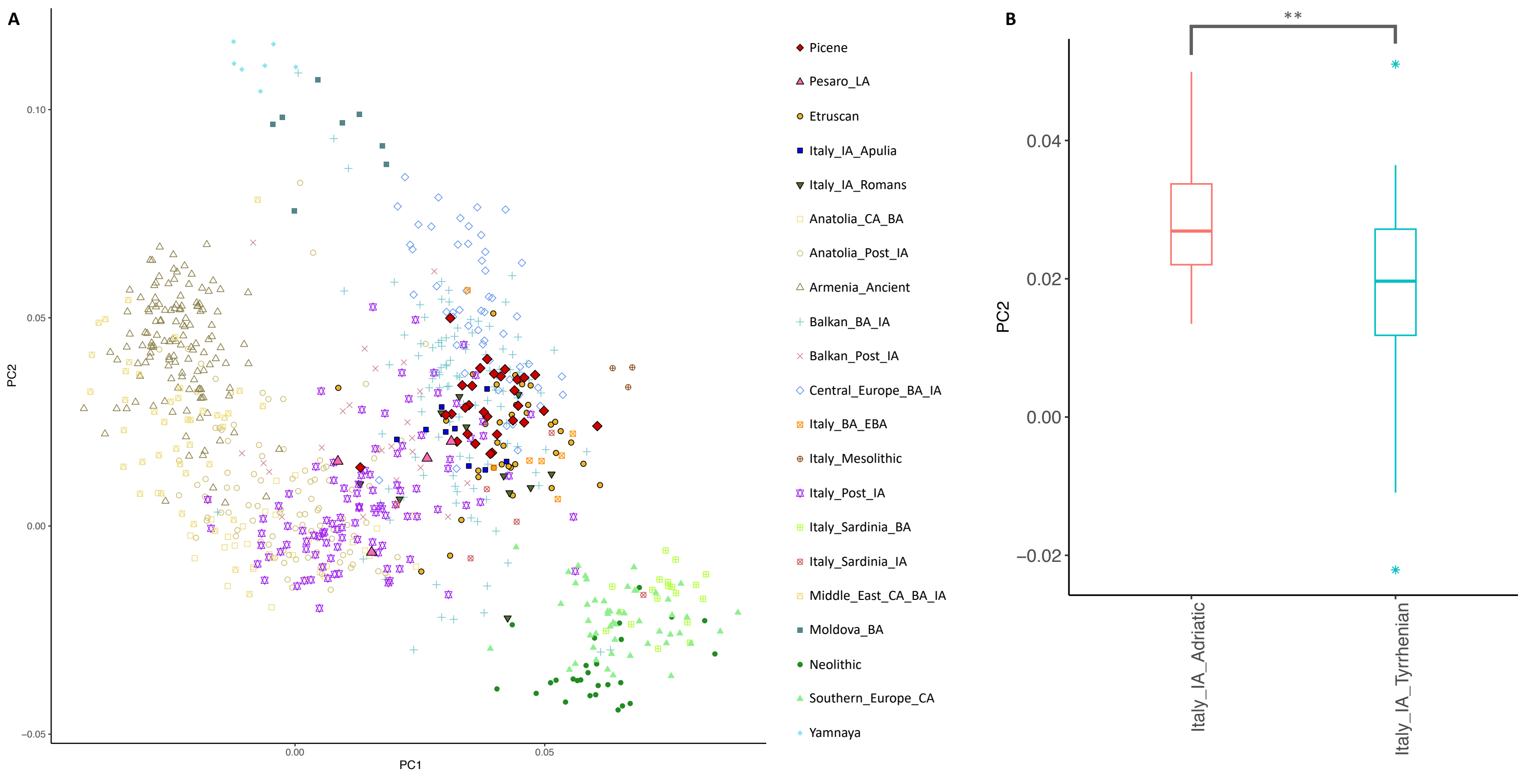


**Figure S7.** A) PCA performed with the total length of shared IBD segments between each pair of individuals included in the analysis (see also Additional file 2: Table S10). Individuals were grouped for relevant periods and geographical origin. Abbreviations: IA=Iron Age, CA=Copper Age, BA=Bronze Age, EBA=Early Bronze Age. B) Boxplots representing the distribution along PC2 of the Italic IA individuals from the Adriatic and the Tyrrhenian side. Colored asterisk in the plot indicate outliers. Significance was calculated with the Mann-Whitney U Test: p-value = 0.00116.

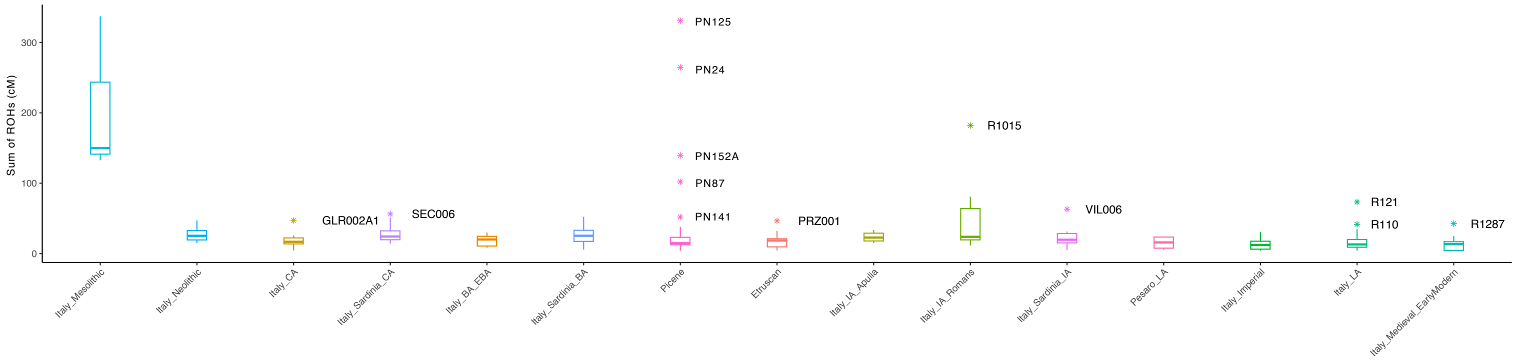


**Figure S8.** Boxplots of the sum of all ROHs > 4cM for individuals of the ancient Italian populations ranging from the Mesolithic to Medieval/Early Modern period. Labels of the outlier individuals are reported.

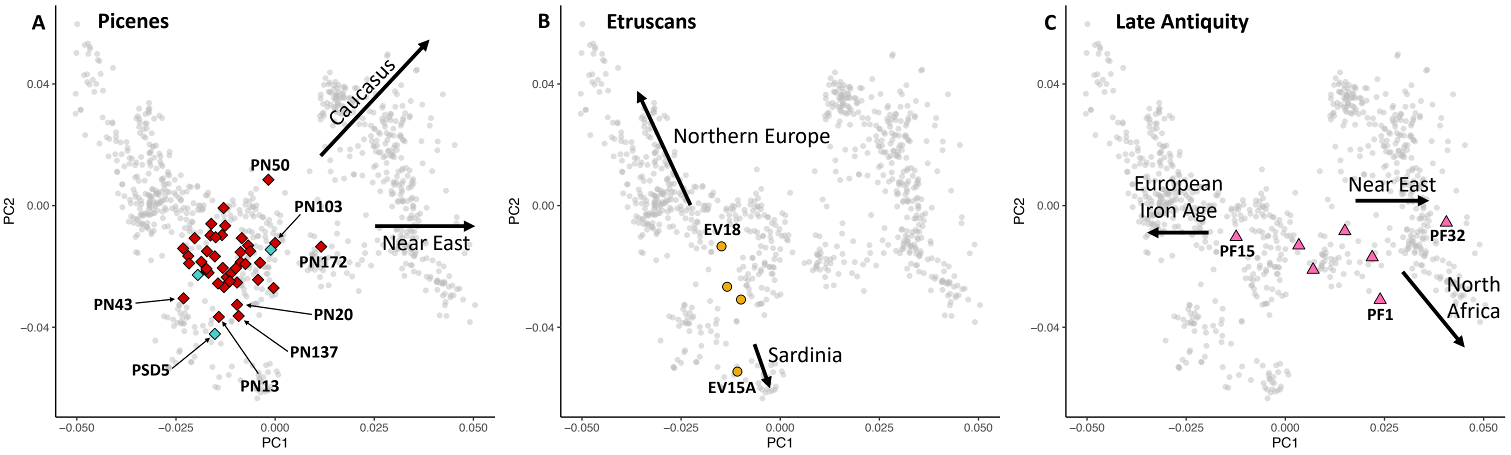


**Figure S9.** PCA highlighting genetic outliers (identified with PCA and/or Admixture analysis) in the groups here analyzed. Modern individuals are in gray. A) Picene individuals. Red squares represent Novilara necropolis, turquoise squares represent Sirolo-Numana necropolis. B) Etruscan individuals from Monteriggioni/Colle di Val d’Elsa. C) Late Antiquity individuals from Pesaro necropolis.

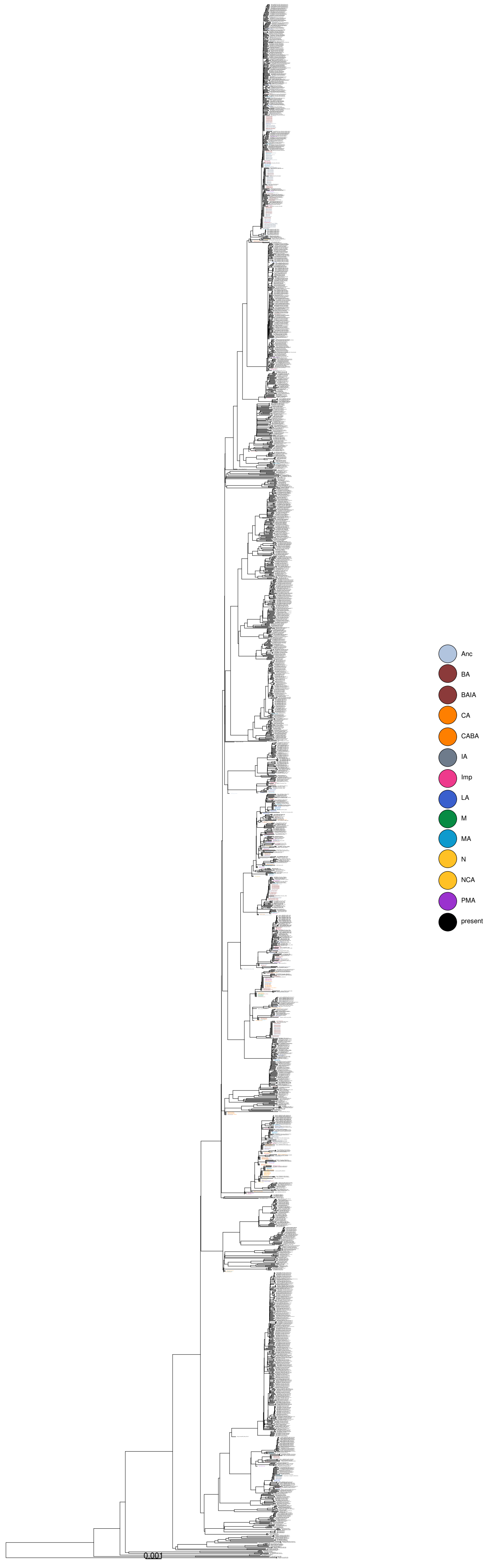


**Figure S10.** Y chromosome phylogenetic tree reconstructed with pathPhynder with both modern and ancient individuals (see Additional file 2: Table S13 for the list of individuals included). Different colors indicate different periods. Abbreviations: Anc=Ancient; BA=Bronze Age; BAIA=Bronze Age/Iron Age; CA=Copper Age; CABA=Copper Age/Bronze Age; IA=Iron Age; Imp=Imperial; LA=Late Antiquity; M=Mesolithic; MA=Middle Ages; N=Neolithic; NCA=Neolithic/Copper Age; PMA=Post Middle Ages.

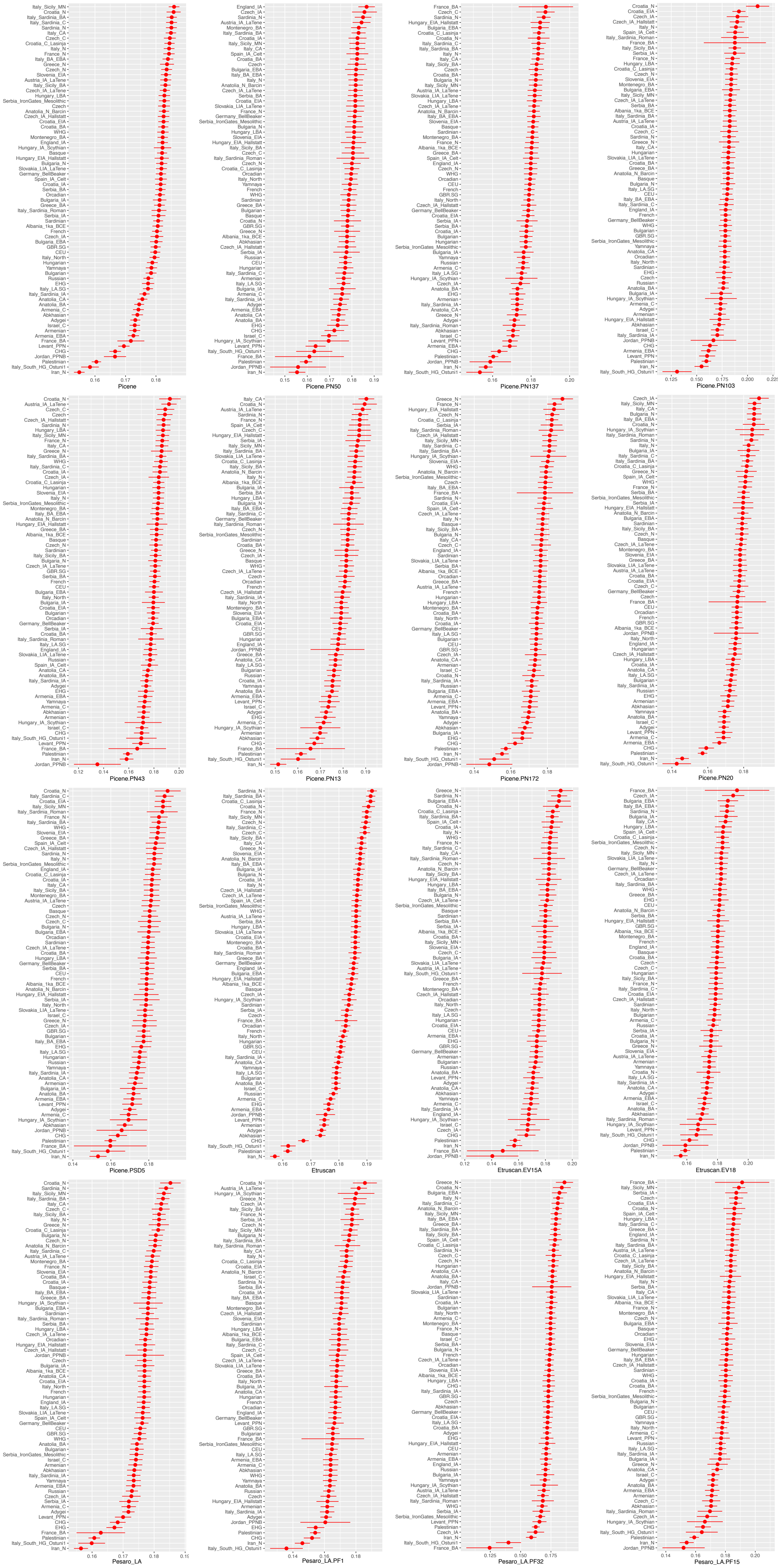


**Figure S11.** f3-outgroup analysis in the form f3(X, Y; YRI.SG) for the Picene, Etruscan and Pesaro_LA general populations and the putative outlier individuals. X population or individual is indicated at the bottom of each plot, Y populations are on the left.
